# An interaction between HP1 and the Chromosomal Passenger Complex Initiates Acentrosomal Spindle Assembly in *Drosophila* oocytes

**DOI:** 10.64898/2026.05.01.722309

**Authors:** Siwen Wu, Ryan Doherty, Manisha Persaud, Keara Greer, Om Patil, Janet K. Jang, Kim McKim

## Abstract

Chromosome segregation fidelity during meiosis is critical for genome integrity, with aneuploidy causing infertility, miscarriages, and congenital anomalies. In the oocytes of many species, spindle assembly occurs in the absence of centrosomes that normally function as microtubule-organizing centers at the poles. Such acentrosomal spindles are believed to pose significant challenges for accurate chromosome segregation compared to centrosomal organized spindles. Previous work in *Drosophila* has shown that the chromosomal passenger complex (CPC) is required for acentrosomal spindle assembly. We found that heterochromatin protein-1 (HP1) plays a critical role in regulating CPC localization and spindle assembly. Furthermore, HP1 moves to the microtubules, where it has roles in building a functional spindle and interacts with the CPC to regulate chromosome biorientation. These results indicate that spindle assembly is mediated by multiple interactions between the CPC, HP1, and the chromosomes, and provide insights into the mechanisms that restricts spindle assembly to the chromosomes in *Drosophila* oocytes.

## Introduction

Accurate chromosome segregation in meiosis requires the assembly of a bipolar spindle^1^. Defects in this process can lead to aneuploidy, infertility, miscarriages, and birth defects^2,3^. Spindle assembly in female oocytes of many species such as humans, mice, *Xenopus*, C. *elegans*, and *Drosophila* are acentrosomal^1,4–7^. For example, *Drosophila* oocytes lack centrosomes and distinct microtubule-organizing centers (MTOC). Instead, after nuclear envelope break down, the chromosomes recruit microtubules and proteins that organize bipolar spindles^6^. The first insights into identifying the molecules required for chromatin-based spindle assembly mechanisms came from work in *Xenopus* egg extracts. DNA-coated beads^7^ or sperm nuclei^8,9^ promote spindle assembly in a process that depends on either RanGTP or the chromosomal passenger complex (CPC)^10,11^. In the absence of the RanGTP gradient, the CPC promotes spindle assembly around sperm nuclei^12^. Furthermore, it has been proposed that the initiation of spindle assembly depends on simultaneous interactions between the CPC, the chromosomes and the microtubules^13^.

Chromosome-mediated spindle assembly in *Drosophila* oocytes depends on the CPC but not the Ran pathway^14,15^. The CPC is composed of four components: the inner centromere protein (INCENP), Borealin, Survivin, and Aurora B kinase^16–18^. Knocking down CPC components INCENP or Survivin in *Drosophila* oocytes results in a failure of spindle and kinetochore assembly during meiosis. Thus, the CPC is required to recruit and organize microtubules to the chromosomes to facilitate acentrosomal spindle formation^19^. Although the CPC has been studied in detail in several systems, the mechanics behind CPC’s recruitment to chromosomes, and its interaction with microtubules to promote spindle assembly are poorly understood. Understanding the mechanism of CPC-mediated spindle assembly should provide insights into how spindle assembly is limited to the vicinity of the chromosomes in oocytes.

Three distinct pathways are known to recruit the CPC to chromosomes. In the first two pathways, the kinases Haspin and Bub1 phosphorylate H3T3 and H2AT120, which respectively recruit CPC components Survivin and Borealin to centromeric chromatin^20,21^. However, knockdown of *Haspin* and *Bub1* does not disrupt spindle assembly in *Drosophila* oocytes, suggesting that these kinases are not required for CPC recruitment^19^. In the third pathway, CPC recruitment to chromosomes is mediated by INCENP and Borealin interactions with heterochromatin protein-1 (HP1, also known as Su(var)205 in *Drosophila*)^18,22–30^. HP1 is a conserved non-histone chromosomal protein associated with heterochromatin^31^. HP1 binds to the methylated K9 residue of histone H3 (H3K9me3), maintaining a transcriptionally repressed state of chromatin^32,33^. Our previous work has shown that the CPC co-localizes with HP1 and H3K9me3 on oocyte chromosomes in experimental situations where the meiotic spindle fails to form, suggesting that HP1 has a role in CPC recruitment to the chromosomes^19^. Additionally, co-localization of HP1 with the CPC on meiotic metaphase spindles has been observed in *Drosophila* oocytes^34^. Based on these observations, we proposed that HP1 plays a role in recruiting the CPC to chromosomes and mediating its transfer to microtubules to initiate acentrosomal spindle assembly.

We have investigated the function of HP1 and its interaction with the CPC in promoting *Drosophila* female meiotic spindle assembly and function. HP1 is required for spindle assembly and chromosome biorientation in oocyte meiosis. HP1 physically interacts with the CPC and recruits it to the chromosomes prior to spindle assembly. We have also directly observed movement of HP1 from the chromosomes to the meiotic spindle. Two domains of INCENP, the SAH and HP1-binding domains, are important for spindle assembly and accurate chromosome segregation in meiosis. These findings reveal a mechanism for spindle formation, based on multiple interactions between the CPC and HP1, and movement of both from the chromosomes to the microtubules.

## Results

### HP1 translocates from heterochromatin to spindle microtubules during meiotic prometaphase I

We first analyzed HP1 localization in *Drosophila* oocyte development. The *Drosophila* ovary typically contains 16 to 20 ovarioles. Each ovariole contains a series of developing oocytes arranged in a linear sequence, progressing through 14 developmental stages that begin in the germarium^35,36^. The germarium contains germline stem cells, mitotic cystoblasts, and oocytes in meiotic zygotene and pachytene. The germarium ends with stage 1 oocytes, at which point the oocytes enter meiotic prophase until stage 12. During this period the chromosomes are condensed into a single spherical structure, or *karyosome.* Entering stage 13, the nuclear envelope breaks down (NEB) to initiate spindle assembly. At stages 13 and 14, the oocyte progresses through prometaphase I and arrests at metaphase I^34^.

Throughout meiotic prophase (stages 1-12), HP1 was localized on the chromosomes while the CPC was absent, indicating the CPC is not loaded until after NEB (Supplemental Figure 1). Thus, HP1 is in position to recruit the CPC to the chromosome prior to spindle assembly. During prometaphase I and metaphase I (stages 13 and 14), *Drosophila* oocytes assemble spindles composed of two types of microtubules: kinetochore fibers (k-fibers) and interpolar fibers^34^. The 5’ ends of K-fibers connect kinetochores to spindle poles, while the 5’ ends of interpolar fibers make antiparallel overlaps in the center of the spindle. The central spindle contains several proteins including the CPC and the kinesin-6 motor protein Subito^37–39^. We observed that HP1 was present on both k-fibers and central spindle fibers (Figure 1A). As the oocytes progress from prometaphase I to metaphase I, the spindle decreases in length^40^. Interestingly, the HP1 localization pattern on spindles was correlated with spindle length. HP1 was enriched in the central spindles when the spindle was relatively long and was more uniformly distributed when the spindle was shorter. These observations suggest a correlation between HP1 distribution and spindle architecture or stage. They also suggest that HP1 can localize on the spindle independently of the CPC.

**Figure 1:**
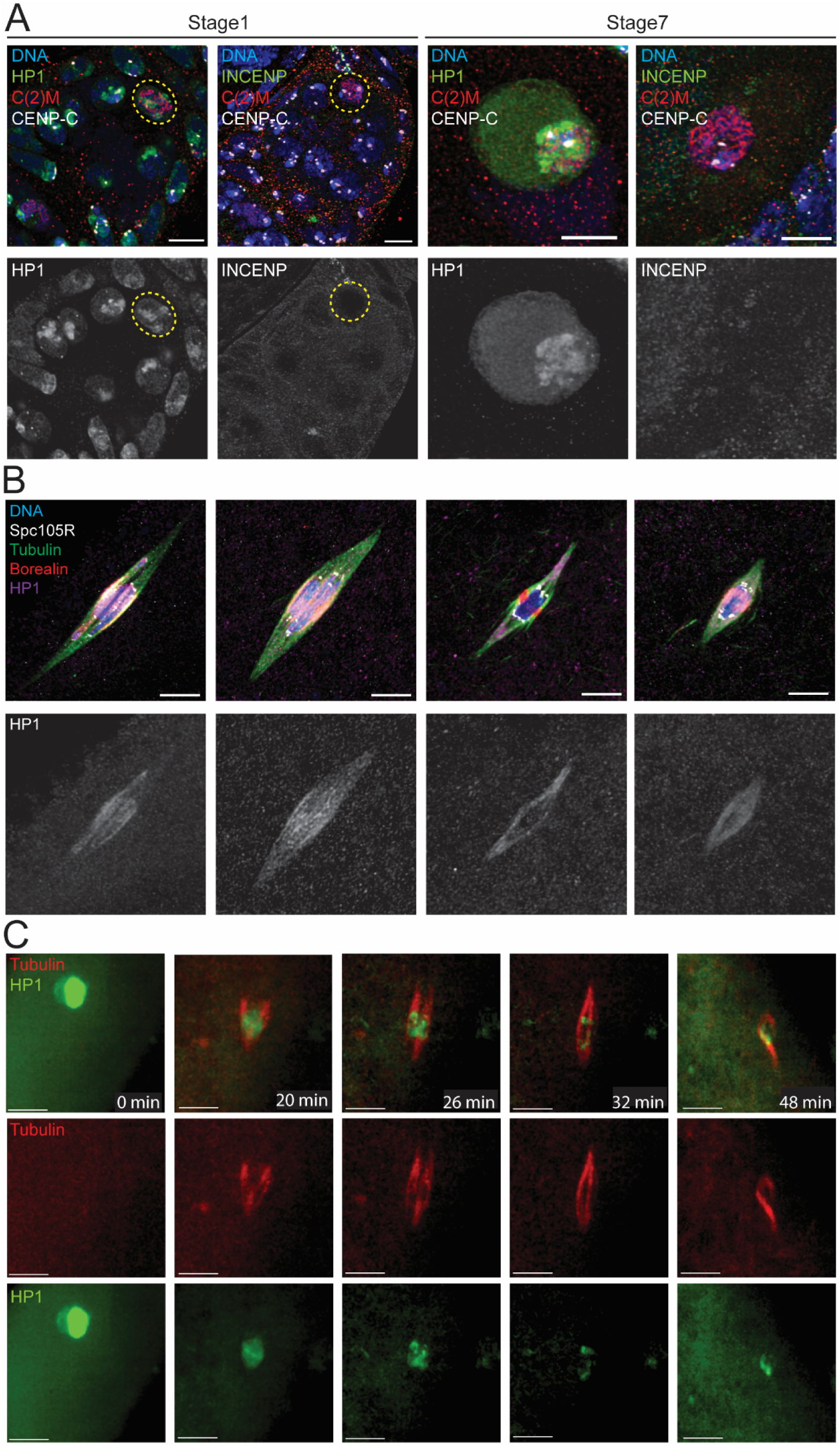
Expression and distribution of HP1 in *Drosophila* female oocytes from meiotic prophase to metaphase I. **(A)** HP1 (green) appeared on chromosomes in prophase I (stage 1 or stage 7) *Drosophila* oocytes. In contrast, the CPC (INCENP, green) was not detected in prophase oocytes, suggesting that CPC is recruited at prometaphase I. C(2)M (red) is a component of the SC in the oocyte and CENP-C (white) is a centromere component. **(B)** Immunofluorescence using antibodies against HP1 (purple), tubulin (green), SPC105R (white) and Borealin (red) indicates that HP1 colocalized with the spindle apparatus at metaphase I. Scale bar is 5 µm. (**C)** Live imaging of a *Drosophila* oocyte entering prometaphase I, using a GFP fusion to HP1and mCherry-Jupiter for the microtubules. HP1 (green) begins in the nucleus. After NEB, the microtubules (red) accumulate around the chromosomes. The time in minutes is shown, and the scale bar is 5µm.

To determine whether HP1 in mitosis shares a similar pattern to that observed in meiosis, we examined HP1 localization in developing embryos. HP1 was absent in early mitotic cycles of the embryo but started localizing in later mitotic cycles 10-11 (Supplemental Figure 1). In later mitotic cycles, HP1 localized on chromosomes during prophase and then translocated to spindle microtubules during metaphase. These observations indicate that HP1 moves from the chromosomes to the microtubules in both the mitotic and meiotic cell divisions of *Drosophila*.

### HP1 moves from the chromosomes to the microtubules after a delay

To directly observe HP1 movement to the spindle, we used a GFP fusion to HP1 at the endogenous locus and mCherry-Jupiter, a microtubule-binding protein, for live imaging (**Error! Reference source not found.**B). Prior to nuclear envelope breakdown, HP1 is localized to the nucleus, with enrichment on the chromosomes. Upon nuclear envelope breakdown, the nucleoplasmic and most of the chromosomal HP1 rapidly dispersed into the cytoplasm. However, a small amount of HP1 remained on the chromosomes while the spindle formed, suggesting a delay in localization of HP1 to the microtubules. Eventually HP1 was observed on the microtubules, suggesting HP1 moves to the spindle men. Although it is possible that spindle HP1 came from the cytoplasm, it is simpler to assume that a fraction of the chromosomal HP1 moves to the spindle.

### HP1 regulates spindle assembly and chromosome biorientation in meiosis

To determine if HP1 plays a role in meiotic spindle assembly, two *Hp1* shRNA transgenes under the control of the UAS/Gal4 system were generated (see Methods, to be referred to as *Hp1^RNAi^*). These shRNAs were expressed by crossing to *matα-GAL4*, which results in RNAi knockdown of HP1 during oocyte prophase development. Expression of either shRNA resulted, 99.9% knockdown of the mRNA, 80% knockdown of the protein and in female sterility (Supplemental Figure 2A, B). We hypothesize that the residual HP1 protein was generated in the early stages of oocyte development before the onset of *matα-GAL4* expression. To test this, we examined HP1 localization throughout meiotic prophase. We observed a mild reduction of HP1 at early prophase (stage 1) and a significant decrease in late prophase (stages 7-10) of *Hp1*^RNAi^ oocytes (Figure 2A-C). These results suggest that some HP1 protein present in *Hp1*^RNAi^ oocytes before the onset of *matα-GAL4* induced expression of the shRNAs persisted until stage 14.

**Figure 2.**
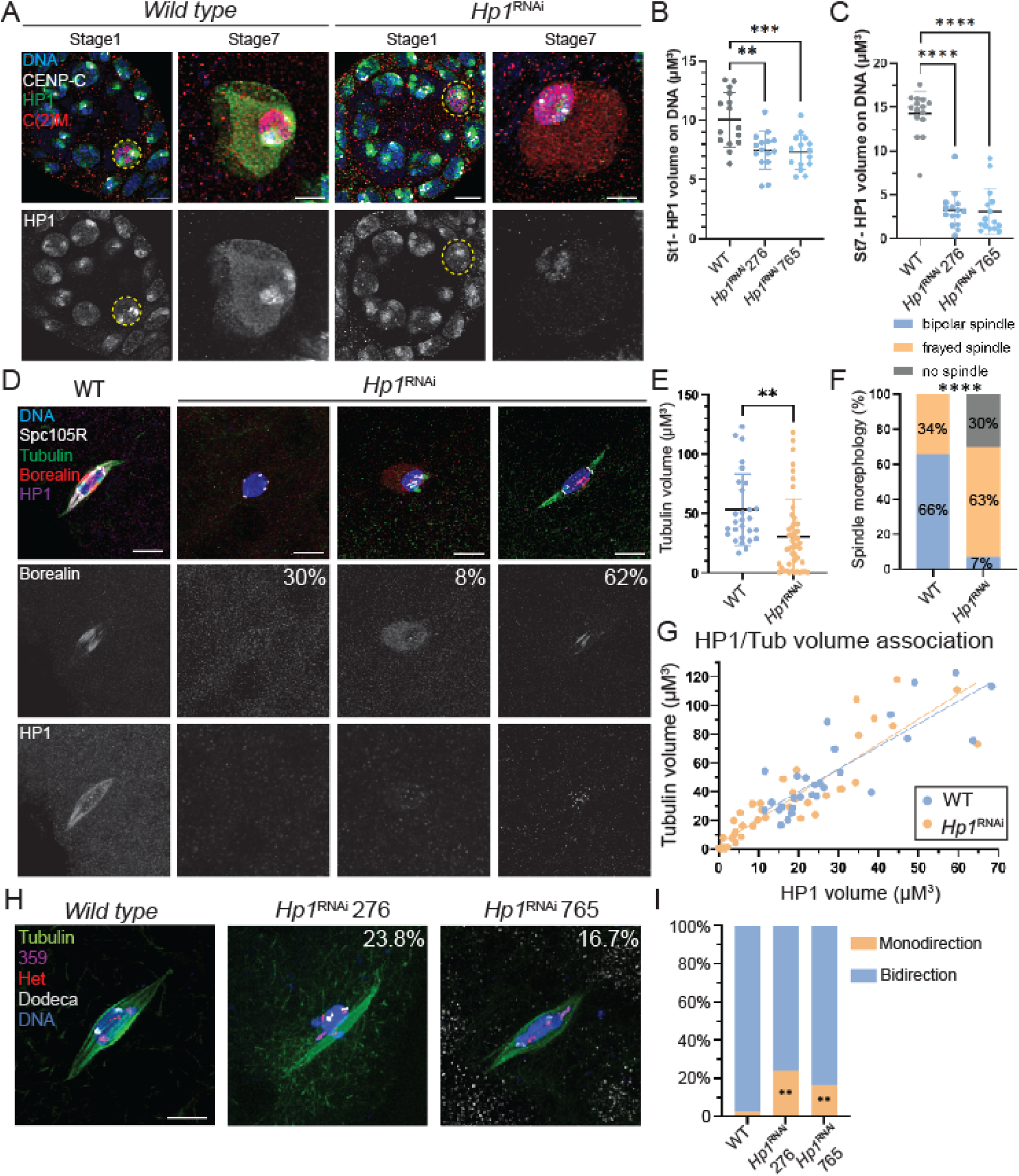
HP1 supports oocyte meiotic development by regulating CPC localization, spindle assembly, and chromosome biorientation. **(A)** Confocal images of wild-type and *Hp1^RNAi^*oocytes at prophase I. Merged images show chromosomes in blue, HP1 in green, CENP-C in white and C(2)M in red. The HP1 channel is also shown separately. HP1 localizes on chromosomes in *wild-type Drosophila* oocytes (highlighted by yellow dashed circles). **(B-C)** Quantification for HP1 volume on DNA in Stage 1 and Stage 7. N = 15 for each group, with ** = p < 0.01, *** = p < 0.001, and **** = p < 0.0001 determined by t-test. **(D)** Confocal images of wild-type and *Hp1^RNAi^* oocytes at metaphase I, with HP1 (purple), CPC (Borealin in red), and spindle (Tubulin in green). 62% of these oocytes showed reduced CPC localization, 30% lacked CPC entirely and 8% displaying delocalized CPC around the karyosome. **(E-G)** Quantitative analysis of spindle volumes (E), spindle morphology (F), and (G) the relationship between HP1 and spindle assembly in *wild-type* (n = 28) versus *Hp1^RNAi^* oocytes (n = 46) (** = p < 0.01 and **** = p < 0.0001 determined by t-test). **(H)** Fluorescence in situ hybridization (FISH) was used to test homologue biorientation in *Hp1^RNAi^*oocytes. Probes against pericentromeric heterochromatin for the X (359 bp repeat, purple), second (Het, red), and third (dodeca, white) chromosomes. **(I)** Quantification of homolog biorientation defects in *wild type* (n = 117), *Hp1^RNAi^* 276 (n = 111) and *Hp1^RNAi^* 765 (n = 102) oocytes (**= P < 0.01, Fisher’s exact test). Oocytes without spindles were excluded from analysis, as bipolarity could not be assessed. Scale bar is 5 µm (all images) and error bars represent 95% confidence intervals.

The effect of HP1 depletion was examined in stage 14 oocytes. In *wild-type* stage 14 oocytes, HP1 was localized to spindle microtubules, including the central spindle (Figure 2D). In *Hp1^RNAi^* oocytes, we observed severe defects in spindle assembly compared with *wild-type*, although there was also a notable variation in spindle volume (Figure 2E). For example, 30% of *Hp1^RNAi^* oocytes failed to assemble spindles, and 63% displayed spindles that were thin and reduced in volume (Figure 2F). In all these oocytes, HP1 was either undetectable or present as puncta on chromosomes in *Hp1^RNAi^*oocytes. The phenotypic variation may have been due to variable amounts of HP1 protein persisting in *Hp1^RNAi^* stage 14 oocytes. Indeed, we observed a correlation between the amount of residual HP1 and the amount of spindle assembly (Figure 2G) in both *wild-type* and *Hp1^RNAi^* oocytes, consistent with a role for HP1 in promoting spindle assembly.

The effect of HP1 depletion on CPC localization was examined using an antibody detecting Borealin. We found that, 62% of *Hp1^RNAi^* oocytes showed reduced CPC localization on the central spindle, with 30% lacking CPC entirely and an additional 8% displaying delocalized CPC around the karyosomes (Figure 2D). These results suggest that HP1 is required for CPC spindle localization and central spindle formation.

One explanation for the variation in *Hp1^RNAi^* oocyte spindle volume is that spindles form that are unstable and degrade over time. For example, incubation of some CPC mutants in modified Robb’s buffer for one hour causes depolymerization of spindle microtubules ^19^. However, *Hp1^RNAi^* oocytes incubated for one hour still formed spindles (Supplemental Figure 2), suggesting that variation in the *Hp1^RNAi^*spindle phenotype was not due to a defect in spindle stability. Thus, our findings demonstrate that HP1 is essential for oocyte spindle assembly.

Proper spindle formation results in the attachment of homologous kinetochores to microtubules from opposite poles, referred to as biorientation at metaphase I. This is essential for accurate separation of homologous chromosomes at anaphase I. To test if *Hp1*^RNAi^ oocytes have defects in biorientation, we performed Fluorescence in situ Hybridization (FISH) assays using probes targeting the pericentromeric regions of chromosomes X, 2, and 3. Homologous chromosomes were considered bi-oriented if two probe signals were oriented towards opposite spindle poles, and mono-oriented if both probes were oriented towards the same spindle pole. Oocytes that failed to form bipolar spindles were excluded from this analysis, as biorientation could not be evaluated without spindle poles. We observed an average of 16.7% rate of mono-oriented chromosomes in the two *Hp1^RNAi^*lines compared to only 3.6% in *wild-type* oocytes, showing that HP1 is required for chromosome biorientation during meiosis I (Figure 2H, I).

### HP1 recruits the CPC to the chromosomes to initiate spindle assembly

To address whether HP1 is involved in recruiting the CPC to chromosomes to initiate spindle assembly, two chromosome binding assays were conducted. First, we used Binuclein 2 (BN2), an inhibitor of Aurora B kinase, which induces microtubule depolymerization^41,42^. In 82.6% of *wild-type* oocytes incubated in BN2 for one hour, spindles were eliminated and CPC component Borealin localized to the chromosomes, indicating movement of the CPC from the microtubules back to the chromosomes (Figure 3A, B). In 100% of *Hp1^RNAi^* oocytes treated with BN2, no spindle formation was observed, along with an 87% reduction in chromosomal HP1 volume, and 55% reduction in Borealin localization on the chromosomes (Figure 3A, B).

**Figure 3.**
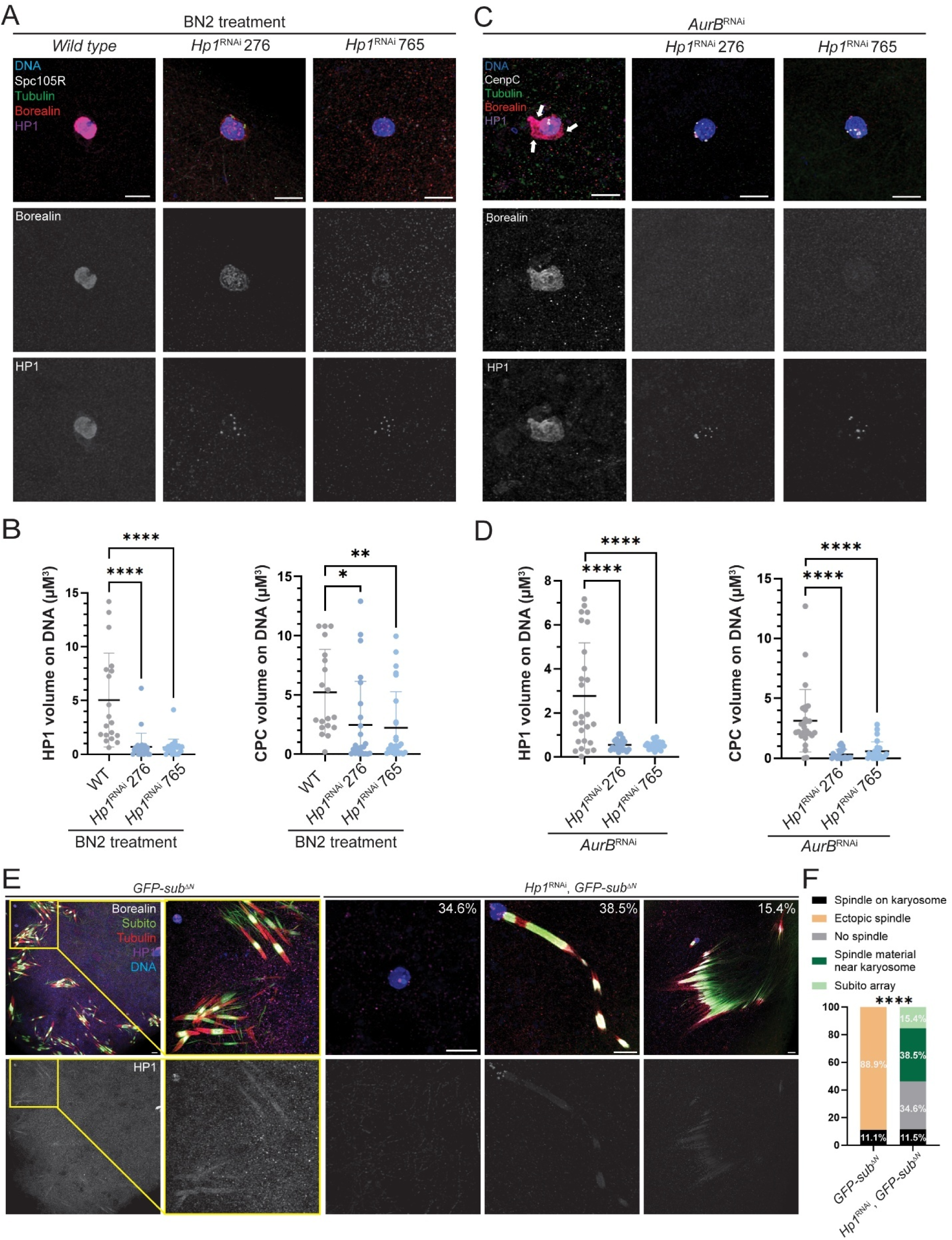
HP1 is required for chromosomal CPC recruitment in meiosis. **(A)** One hour incubation of oocytes in the Aurora B inhibitor BN2 causes loss of the meiotic spindle and localization of CPC component Borealin (red) and HP1 (purple) to the chromosomes. Also shown are SPC105R (white), tubulin (green) and DNA (blue). **(B)** Quantitative analysis of *Hp1^RNAi-276^* (n = 25) and *Hp1^RNAi-765^* (n = 27) oocytes compared with *wild type* (n = 19) (t-test: *= P < 0.05, ** = P < 0.01, **** = P <0.0001). **(C)** Aurora B (AurB) was depleted via RNAi to test CPC (Borealin, red) and HP1 (purple) localization prior to spindle assembly, with CENP-C in white. The white arrows show the concentration of HP1 and the CPC not colocalizing with the spindle or chromosomes. **(D)** Quantification of HP1 and CPC volume on chromosomes in *AurB^RNAi^* (n = 27), *Hp1^RNAi-276^*; *AurB^RNAi^* (n = 22), and *Hp1^RNAi-765^*; *AurB^RNAi^* (n = 25) oocytes (t-test: **** = P <0.0001). Error bars represent 95% confidence intervals. **(E)** *Drosophila* kinesin-like protein Subito lacking its N-terminal domain (*GFP-sub*^Δ*N*^) induces ectopic spindle formation in stage 14 oocytes^50^. DNA is blue, Borealin is white, Subito is green, HP1 is purple and tubulin is red. **(F)** Quantification of ectopic spindle assembly based on 18 *GFP-sub*^Δ*N*^ oocytes and 26 *Hp1^RNAi^ GFP-sub*^Δ*N*^oocytes using (Fisher’s exact test). **** = P <0.0001. Scale bar, 5 µm (all images).

In the second chromosome binding assay, we knocked down Aurora B by RNAi (*AurB^RNAi^*) to arrest oocytes before spindle assembly. In *AurB^RNAi^* oocytes, both HP1 and the Borealin colocalized on the chromosomes (Figure 3C-D). However, in the double depletion *AurB^RNAi^*, *Hp1^RNAi^* oocytes, chromosomal HP1 volume was reduced by 80.9%, and Borealin localization on the chromosomes was reduced by 85.7%, suggesting that HP1 is essential for recruiting the CPC to chromosomes prior to spindle assembly. These results indicate that HP1 is required for CPC recruitment to chromosomes.

An important difference between BN2-treated and *AurB^RNAi^*oocytes is whether the spindle forms first and then depolymerized (as with BN2), or if it never forms (as in *AurB^RNAi^*). With BN2 treatment, CPC components that had localized to spindles returned to the chromosomes upon spindle disassembly. In *AurB^RNAi^*, the CPC localizes to the chromosomes and no spindle assembly occurs. This distinction may cause the subtle differences in the results observed in the two conditions. Both experiments do show, however, that HP1 is required for CPC localization to the chromosomes. However, the second (*AurB^RNAi^*) assay may better represent the effect of HP1 on initial CPC recruitment.

### HP1 is required for chromosome-independent spindle assembly

The above experiments show that HP1 is required to recruit the CPC to the chromosomes. To determine if HP1 has a function in spindle assembly while localized to the microtubules, we used oocytes where spindles are assembled without chromosomes. Subito is a kinesin-6 required for bundling of central spindle microtubules and spindle bipolarity in *Drosophila* oocytes^37,43^. A *sub* mutant containing a deletion of the N-terminal domain (*GFP-sub*^Δ*N*^) induces ectopic spindle formation by organizing and bundling antiparallel microtubules that are not associated with chromosomes (Figure 3E). These ectopic spindles depend on the CPC, suggesting that Subito can recruit the CPC and promote spindle formation^19^. If HP1 is required for the CPC to promote spindle assembly, then these spindles should contain HP1 protein, and HP1 depletion should prevent ectopic spindle assembly. In controls, 88.9% of *GFP-sub*^Δ*N*^ oocytes formed ectopic spindles and HP1 localized to these ectopic spindles. In contrast, *Hp1^RNAi^ GFP-sub*^Δ*N*^ oocytes failed to produce ectopic spindles (Figure 3E).

Instead, 34.6% of oocytes showed complete absence of spindle formation, similar to the frequency of *Hp1^RNAi^* oocytes that lack a spindle (Figure 2F). Novel spindle assembly phenotypes were also observed, with either linear (38.5%) or parallel arrays (15.4%) of Subito associated with the CPC and tubulin adjacent to the karyosome. The absence of ectopic spindles suggests that, in addition to chromosome recruitment, HP1 has a role in spindle assembly while located on the microtubules, possibly involving an interaction with the CPC.

### HP1 is required for mitosis in *Drosophila* embryos

To determine whether the loss of HP1 in oocytes affects subsequent embryonic development, we examined embryos from *Hp1^RNAi^*mothers. Following a 1-hour collection, embryos from wild-type mothers were observed at a range of developmental stages, including early mitotic divisions and later stages in which nuclei migrated to the surface and were evenly distributed that contained HP1 on the spindles (Figure 4A).

**Figure 4.**
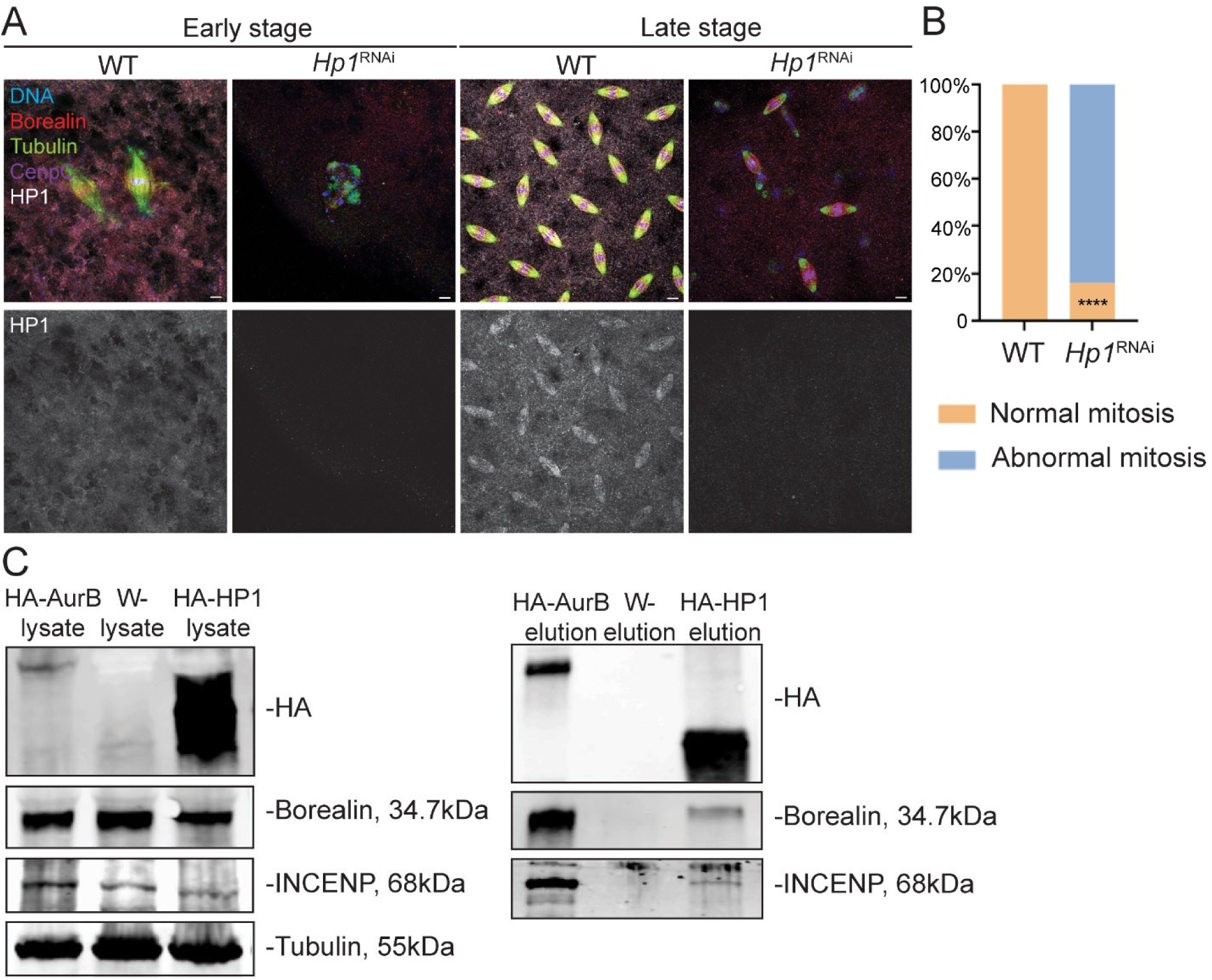
HP1 is required for mitotic progression in *Drosophila* embryos and interacts with the CPC. **(A)** *Drosophila* embryos were collected at 2-hour intervals to test the function of HP1 in mitosis. DNA is shown in red, Borealin in red, HP1 in white, CENP-C in purple, and microtubules in green. Scale bar is 5 µm. **(B)** The frequency of mitotic abnormalities in 34 *wild type* and 25 *Hp1^RNAi^* embryos. (**** = P <0.0001, Fisher’s exact test). **(C)** Co-immunoprecipitation (Co-IP) was conducted to identify potential binding partners of HP1. Wild-type embryos served as a negative control, while *HA-AurB* expressing embryos were used as a positive control. Lysates were prepared and approximately 2 mg of total protein per sample was used for each experiment. In the wild-type control (*w^-^*, middle column), INCENP and Borealin were detected in the input but not in the elution fraction. Positive control (*HA-AurB*, left column) confirmed that Aurora B associated with both INCENP and Borealin. HP1 was found to co-IP with INCENP and Borealin (right column). Tubulin was used as a loading control.

Under the same conditions, we observed two types of defects in *Hp1^RNAi^*embryos (Figure 4B). First, 68% (n=25) of embryos were at an early stage of development with abnormal spindles, indicating an early mitotic and developmental arrest. Second, the mitotic nuclei had migrated to the surface in 16% of embryos, but some nuclei appeared to have sunk back toward the center, probably due to errors in the mitotic divisions. The remaining 16% of embryos underwent normal mitotic divisions but ultimately must have failed to develop, given the complete embryonic lethality. These findings suggest that maternal HP1 expression in the oocyte is required for mitosis during early embryogenesis, consistent with previous findings showing that HP1 deficiency leads to cell division defects in *Drosophila* embryos^44,45^.

### HP1 interacts with CPC components INCENP and Borealin

To determine if there is a physical interaction between the CPC and HP1 in vivo, co-IP experiments were conducted. Due to the low spindle-to-cytoplasm ratio in metaphase I oocytes, and because the localization pattern of the CPC is similar in oocytes and embryos, we used embryos with mitotically dividing nuclei to investigate CPC-HP1 interactions with co-IP experiments. Using embryos expressing HA-tagged Aurora B as a positive control, we found that both INCENP and Borealin were detected in the input and HA-AuroraB pull-down fractions, confirming these protein interactions (Figure 4C). Importantly, HA-HP1 co-immunoprecipitated with both INCENP and Borealin, indicating that HP1 interacts with CPC components in embryonic mitotic cells (Figure 4C). INCENP and Borealin were detected in the input but were absent in the HA pull-down from wild-type embryos that lacked the HA tag, confirming the specificity of the Co-IP. These results indicate that HP1 physically interacts with the CPC.

### The STD and SAH domains of INCENP is not required for spindle assembly

To investigate the interactions between the CPC, HP1 and microtubules required for meiosis I spindle assembly and function, we generated and analyzed a series of *Incenp* mutants (Figure 5A). INCENP has the C-terminal INbox domain that recruits Aurora B kinase, and several other domains that target the CPC to a variety of locations. Its N-terminal domain (BS) binds to Borealin and Survivin. The STD domain in human INCENP has been shown to interact with the homolog of kinesin-6 Subito, Mitotic Kinesin-Like Protein 2 (MKLP-2/Kif20a)^17^. In addition to SAH, the STD domain have been proposed to interact with microtubules^22,46^. A phospho-regulatory domain (PRD) located upstream of the SAH domain contains six potential CDK1 phosphorylation sites in human and Xenopus^47^, while four potential CDK1 sites were identified in the corresponding PRD region in *Drosophila*. Between STD and PRD is a domain containing HP1 binding sites^24,29^. The expression of the mutated *Incenp* transgenes in oocytes was verified by Western blot (Supplemental Figure 3). Most of the mutants failed to rescue the sterility phenotype of *Incenp^RNAi^*, and were sterile in the presence of wild-type *Incenp,* indicating they had a dominant negative phenotype (Table 1).

**Figure 5.**
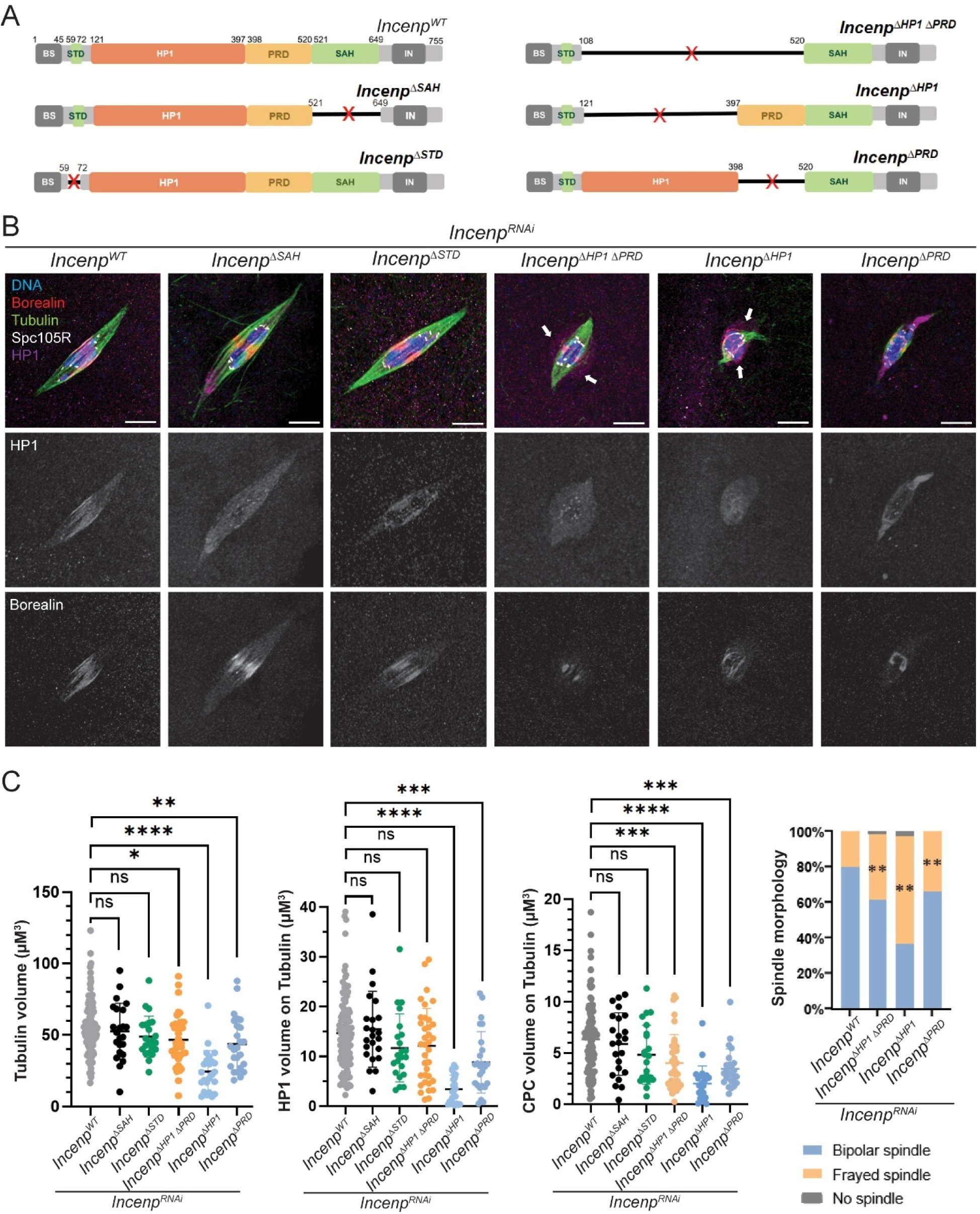
HP1-INCENP interaction is not sufficient for spindle formation, but is required for building a functional central spindle in meiosis. A schematic of *Drosophila* INCENP deletion constructs designed to delete each functional domain. Microtubule (MT) binding domains STD (59-72) and SAH (single α-helix, 521-649) are deleted in *Incenp*^Δ*STD*^ and *Incenp*^Δ*SAH*^ transgenes, respectively. The potential HP1-binding region (amino acids 121–397) is deleted in *Incenp*^Δ*HP1ex*^. The phospho-regulatory domain (PRD, 398-520), containing CDK phosphorylation sites upstream of the SAH domain^51^, was deleted in *Incenp*^Δ*PRD*^. **(B)** HP1 (purple), CPC/Borealin (red), Tubulin (green) and SPC105R (white) were examined in oocytes expressing RNAi-resistant *Incenp* transgenes in an *Incenp^RNAi^* background. The white arrows show the concentration of HP1 not colocalizing with the spindle or chromosomes. Scale bar, 5 µm (all images). **(C)** Quantification of data in *Incenp^WT^* (n = 108), *Incenp*^Δ*SAH*^ (n = 23), *Incenp*^Δ*STD*^ (n = 22), *Incenp*^Δ*HP1*^ ^Δ*PRD*^ (n = 36), *Incenp*^Δ*HP1ex*^ (n = 24), and *Incenp*^Δ*PRD*^ (n = 25) oocytes (all in *Incenp^RNAi^* background). Error bars represent 95% confidence intervals and * = P < 0.05, ** = P < 0.01, *** = P < 0.001, **** = P <0.0001 by t-test.

**Table 1.** Summary of transgenes, fertility and nondisjunction.

| Genotype | Deletion | Fertility/NDJ<sup>a</sup> |  |
| --- | --- | --- | --- |
|  |  | In WT oocytes | In *Incenp* RNAi oocytes |
| *Incenp*<sup>WT</sup> |  |  | +++/0.5% (*n* = 920) |
| *Incenp*<sup>ΔSTD</sup> | 59-78 | +++/0% (*n* = 310) | ++/0.2% (*n* = 600) |
| *Incenp*<sup>Δm</sup> | 59-649 | Sterile | Sterile |
| *Incenp*<sup>ΔHP1<sup>ex</sup></sup> | 108-397 | Sterile | Sterile |
| *Incenp*<sup>ΔPRD</sup> | 398-520 | Sterile | Sterile |
| *Incenp*<sup>ΔSAH</sup> | 521-649 | +/0% (*n* = 54) | Sterile |
| *Incenp*<sup>ΔHP1ΔPRD</sup> | 108-520 | Sterile | Sterile |
| *Incenp*<sup>ΔHP1ΔST</sup> | 59-78, 108-397 | Sterile | Sterile |
| *Incenp*<sup>ΔHP1ΔSAH</sup> | 108-397, 521-649 | Sterile | Sterile |
| *Incenp*<sup>ΔSTDΔSAH</sup> | 59-78, 521-649 | Sterile | Sterile |

NDJ = 2XNDJ/total progeny.
<sup>a</sup>Females were crossed to *y Hw w/BSY* males in vials. Fertility is based on the number
of progeny per vial: +++ = 20–50 per vial; ++ = 10–20 per vial; and + = 1–10 per vial;
sterile = no progeny.

We examined the effect of these mutants on spindle assembly, and HP1 and Borealin localization in a *Incenp^RNAi^* background. Previous studies in vertebrate cells have used a minimal INCENP fragment containing the first 54-80 amino acids to target the CPC to chromosomes^26,48^. In contrast, a similar mutant lacking the middle region between the BS and the INbox domains failed to localize to chromosomes or form spindles (Supplemental Figure 4). Therefore, at least one domain among STD, HP1, PRD and SAH must contribute to chromosome localization and spindle assembly. To determine if the STD and SAH domains are required for microtubule assembly, both domains were deleted. Surprisingly, deletion of both microtubule-binding domains (DSTDDSAH) did not impair spindle formation (Supplemental Figure 4). These findings suggest that the central region of INCENP containing the HP1 and PRD domains may be sufficient for meiotic spindle assembly.

### The HP1 domain of INCENP is required for spindle organization but not assembly

We assessed spindle morphology in mutants lacking the HP1 and PRD domains and compared to mutants lacking the STD and SAH domains. All genotypes were able to form spindles at metaphase I (Figure 5B). However, spindle volume and CPC (Borealin) volume on spindles was significantly reduced in *Incenp*^Δ*HP1ex*^ and *Incenp*^Δ*PRD*^, but were unaffected in *Incenp*^Δ*STD*^and *Incenp*^Δ*SAH*^ (Figure 5C). Consistently, this reduction in Borealin levels correlated with decreased overall INCENP localization in these mutants (Supplemental Figure 5A). Despite these defects, the majority of oocytes still formed spindles, indicating that no single domain of INCENP is required for spindle assembly. In contrast, because *Incenp*^Δ*HP1ex*^ displayed the most severe defects, an interaction between HP1 and INCENP may support proper spindle structure and function during metaphase I.

We examined HP1 localization in each mutant to assess the interaction between HP1 and INCENP during spindle assembly (Figure 5B, C). As described above, HP1 normally localizes to metaphase I spindles, with enrichment in the central spindle. Quantitative analysis showed a significant reduction in spindle-associated HP1 volume in *Incenp*^Δ*HP1ex*^and *Incenp*^Δ*PRD*^ oocytes but not *Incenp*^Δ*SAH*^ and *Incenp*^Δ*STD*^. These results suggest that the HP1-binding domain in INCENP is required for HP1 recruitment to meiotic spindles.

In *Incenp*^Δ*HP1ex*^ (n = 13/43) and *Incenp*^Δ*HP1*Δ*PRD*^ (n = 21/42) oocytes, we observed HP1 localization extending beyond the microtubules and chromosomes (Figure 5B). Similarly, non-chromosomal material and non-spindle concentration of HP1 was observed in *AurB^RNAi^* oocytes (n = 15/45, Figure 3C). Interestingly, in 6/15 *AurB^RNAi^* oocytes the non-chromosomal HP1 concentrated together with CPC. These observations may be similar to observations in previous studies showing that HP1 and the CPC can undergo liquid–liquid phase separation (LLPS) during mitosis^26,49^. We speculate that the HP1 aggregation observed near chromosomes in *Incenp*^Δ*HP1ex*^and *AurB^RNAi^* oocytes may be associated with this LLPS behavior. The fact that HP1 has this activity without the CPC (eg. *Incenp*^Δ*HP1ex*^ oocytes) suggests HP1 is driving this activity.

To directly examine central spindle formation, we used localization of Subito. In wild-type oocytes, Subito colocalizes with the CPC in the central spindle microtubules. In both *Incenp*^Δ*HP1ex*^ and *Incenp*^Δ*PRD*^ mutants, Subito localization was significantly reduced, with the greatest reduction seen in *Incenp*^Δ*HP1ex*^ (Supplemental Figure 5B-C). Thus, loss of the HP1 interaction domain impairs Subito recruitment, resulting in reduced central spindle structure. In conclusion, while the HP1 domain of INCENP is not required for CPC recruitment to the chromosomes, it is required for CPC localization and assembly of the central spindle.

Subito has a MT-binding domain within its C-terminal domain and interacts with the CPC^50^ and a HP1 interaction site^19^. Thus, it is possible that a Subito – CPC or Subito – HP1 interactions could promote spindle assembly. To test whether HP1 and Subito function cooperatively in recruiting the CPC to the microtubules, we combined *subito^RNAi^* with *Incenp* mutants (Supplemental Figure 5D). In *subito^RNAi^* oocytes, multipolar or monopolar spindles form and the CPC localizes on all the microtubules, consistent with previous data that the localization of the CPC to the central spindle depends on Subito^19,39^. In *subito^RNAi^*, *Incenp*^Δ*HP1ex*^ oocytes, Borealin was observed on microtubules. However, Tubulin was reduced and disorganized and HP1 was delocalized in 17 out of 30 *Incenp*^Δ*HP1ex*^ oocytes. These results suggest that Subito may interact with and stabilize both HP1 and the CPC on the spindle.

### Evidence for interactions between domains regulating spindle assembly

Deletion of the HP1 domain of INCENP resulted in reduced central spindle assembly, while deletion of the entire middle region of INCENP abolished it entirely. This led us to hypothesize that spindle assembly may depend on contributions from at least two domains within INCENP. To test if multiple INCENP domains are required for spindle assembly, we generated double-deletion *Incenp* mutants lacking the HP1-binding domain in combination with either the MT-binding STD or SAH domains, or the PRD domain (Figure 6A). *Incenp*^Δ*STD*Δ*HP1*^ and *Incenp*^Δ*HP1*Δ*PRD*^oocytes did not have a more severe spindle phenotype compared to *Incenp*^Δ*HP1ex*^. In contrast, *Incenp*^Δ*HP1*Δ*SAH*^ oocytes did have a more severe phenotype, failing to assemble metaphase I spindles (Figure 6B, C). Consistent with these phenotypes, both CPC and HP1 localization were absent in *Incenp*^Δ*HP1*Δ*SAH*^oocytes. These findings suggest that the HP1 and SAH domains both contribute to spindle assembly in *Drosophila* oocytes.

**Figure 6.**
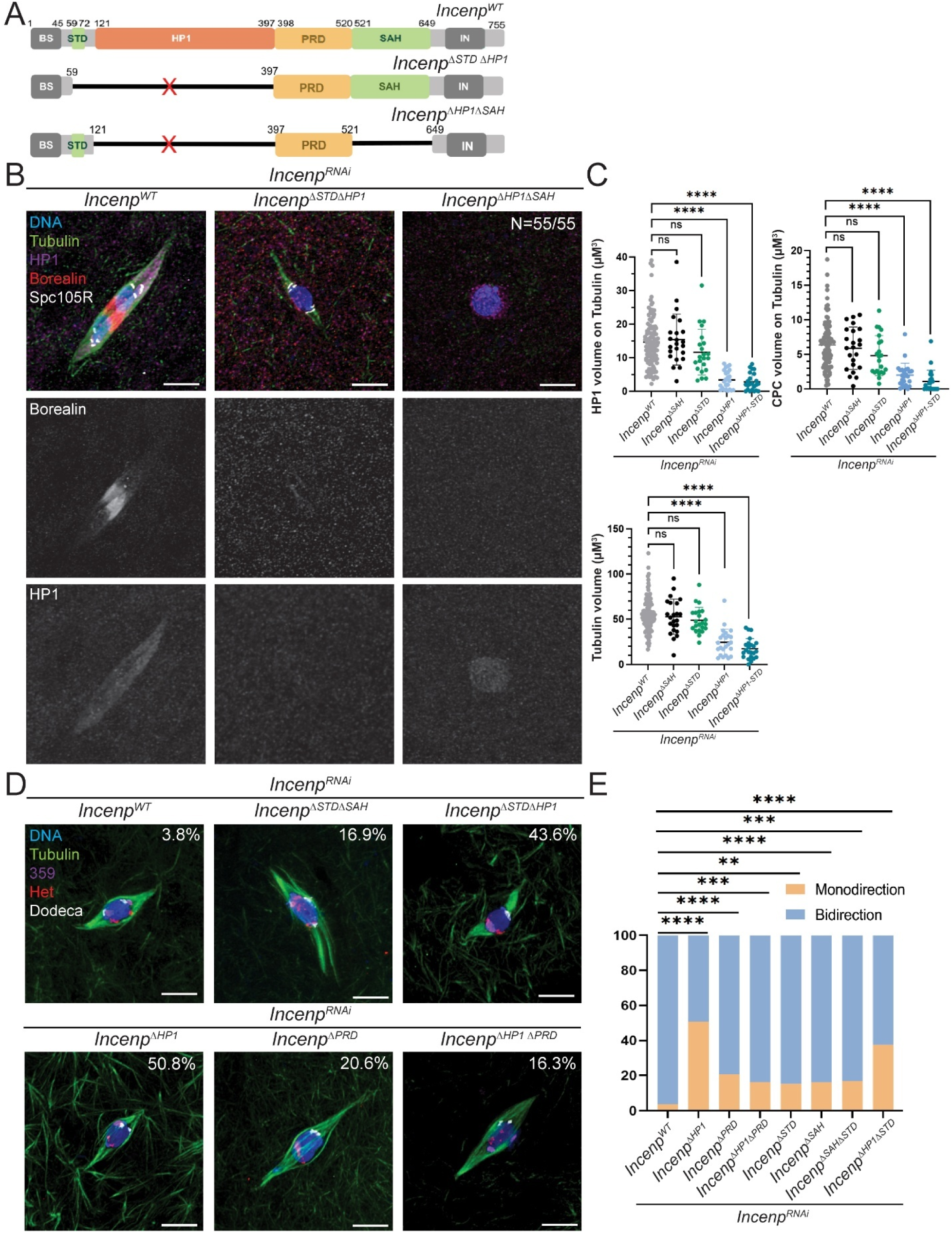
The HP1 and SAH domains of INCENP contribute additively to spindle assembly and chromosome biorientation during meiosis. **(A)** Schematic representation of *Drosophila* INCENP deletion constructs of the HP1-binding and MT-binding domains. The mutant *Incenp*^Δ*STD*Δ*HP1*^ lacks both the STD and HP1 binding domains, *Incenp*^Δ*HP1*Δ*SAH*^ lacks the HP1 and SAH domains, and *Incenp*^Δ*HP1*Δ*PRD*^ lacks both PRD and HP1 binding domains. **(B)** Representative images of stage 14 oocytes stained for HP1 (purple), CPC/Borealin (red), SPC105R (white) and tubulin (green) to assess spindle morphology with double deletion transgenes. **(C)** Quantification analysis of HP1 and CPC volume on spindles (upper panel) and Tubulin volume (lower panel). Data from *Incenp^WT^* (n = 108), *Incenp*^Δ*SAH*^(n = 23), *Incenp*^Δ*STD*^ (n = 22), *Incenp*^Δ*HP1ex*^(n = 24), and *Incenp*^Δ*HP1*^ ^Δ*STD*^(n = 23) oocytes (all in *Incenp^RNAi^* background). The *Incenp*^Δ*HP1*Δ*SAH*^mutant is not shown on the graphs because all oocytes lacked a spindle (n=55). Error bars indicate 95% confidence intervals and significance was analyzed by t-test (**** = P < 0.0001). **(D)** FISH experiments were performed on *Incenp* mutants to examine homologous chromosome biorientation. Mono-orientation rates are indicated at the top right of each representative image. **(E)** Quantification of the frequency of biorientation defects in *Incenp^WT^* (n = 123), *Incenp*^Δ*SAH*^ (n = 60), *Incenp*^Δ*STD*^ (n = 60), *Incenp*^Δ*HP1*^ ^Δ*PRD*^(n = 135), *Incenp*^Δ*HP1ex*^ (n = 132), *Incenp*^Δ*PRD*^(n = 102), and *Incenp*^Δ*HP1*^ ^Δ*STD*^(n = 150) oocytes, all in an *Incenp^RNAi^* background. FISH data analyzed using Fisher’s exact test, with ** = P < 0.01, *** = P < 0.001, and **** = P < 0.0001. Scale bar, 5 µm (all images).

The *Incenp*^Δ*HP1ex*^ mutant consistently had a more severe phenotype than *Incenp*^Δ*HP1*Δ*PRD*^, including defective spindle volume, reduced CPC and HP1 on DNA, and chromosome biorientation defects (Figure 5B-C, Figure 6D-E). These results can be explained if the PRD domain has a negative regulatory effect on spindle assembly, or more specifically, in CPC binding to microtubule as proposed in vertebrate cells^51^.

### HP1-INCENP interaction is required for chromosome bi-orientation

To determine which domains of INCENP are required for chromosome alignment, FISH analyses were performed across all *Incenp* mutants (Figure 6D). We hypothesized that disruption of the HP1–INCENP interaction, as well as the INCENP–MT interaction, would impair chromosome bi-orientation. Indeed, meiotic metaphase I oocytes expressing *Incenp* mutants lacking MT-binding domains (*Incenp*^Δ*STD*^, *Incenp*^Δ*SAH*^ and *Incenp*^Δ*STD*Δ*SAH*^) revealed a significant increase in chromosome segregation errors compared to controls (Figure 6E). Similarly, deletion of the HP1-binding domain (*Incenp*^Δ*HP1ex*^) or the PRD domain (*Incenp*^Δ*PRD*^), or both (*Incenp*^Δ*HP1*Δ*PRD*^), also led to significantly elevated biorientation defects. Among all mutants, *Incenp*^Δ*HP1ex*^ and *Incenp*^Δ*STD*Δ*HP1*^ showed the highest frequency of bi-orientation errors, suggesting that the HP1–INCENP interaction plays an important role in promoting chromosome alignment, potentially more critical than the MT-binding domains of INCENP. Interestingly, *Incenp*^Δ*HP1ex*^ had significantly more defects in chromosome biorientation than *Incenp*^Δ*HP1*Δ*PRD*^. This is consistent with the defects observed in the CPC and spindle structures, suggesting that the PRD domain negatively regulates interactions between INCENP and the microtubules.

### The SAH and HP1 domains of INCENP have different roles in CPC recruitment

To determine whether the interaction between HP1 and INCENP plays a role in recruiting the CPC to chromosomes, we employed the chromosome binding assays used with *Hp1^RNAi^* oocytes. In *wild-type* oocytes treated with BN2, spindle depolymerization caused CPC and HP1 localization to the chromosomes (Figure 7A), indicating that both HP1 and the CPC translocate from spindles back to chromosomes upon spindle disassembly. BN2 treatment of the *Incenp*^Δ*SAH*^ mutant showed no significant defect in CPC recruitment to the chromosomes, even though HP1 was not changed. In contrast, mutants lacking the HP1-binding domain, including *Incenp*^Δ*HP1ex*^, *Incenp*^Δ*HP1*Δ*SAH*^, *Incenp*^Δ*HP1*Δ*STD*^, and *Incenp*^Δ*m*^, showed reduced CPC and HP1 localization on the chromosomes (Figure 7A-B). These results suggest that the HP1 domain in INCENP contributes to CPC recruitment to the chromosomes, although this could be related to reduced HP1.

**Figure 7.**
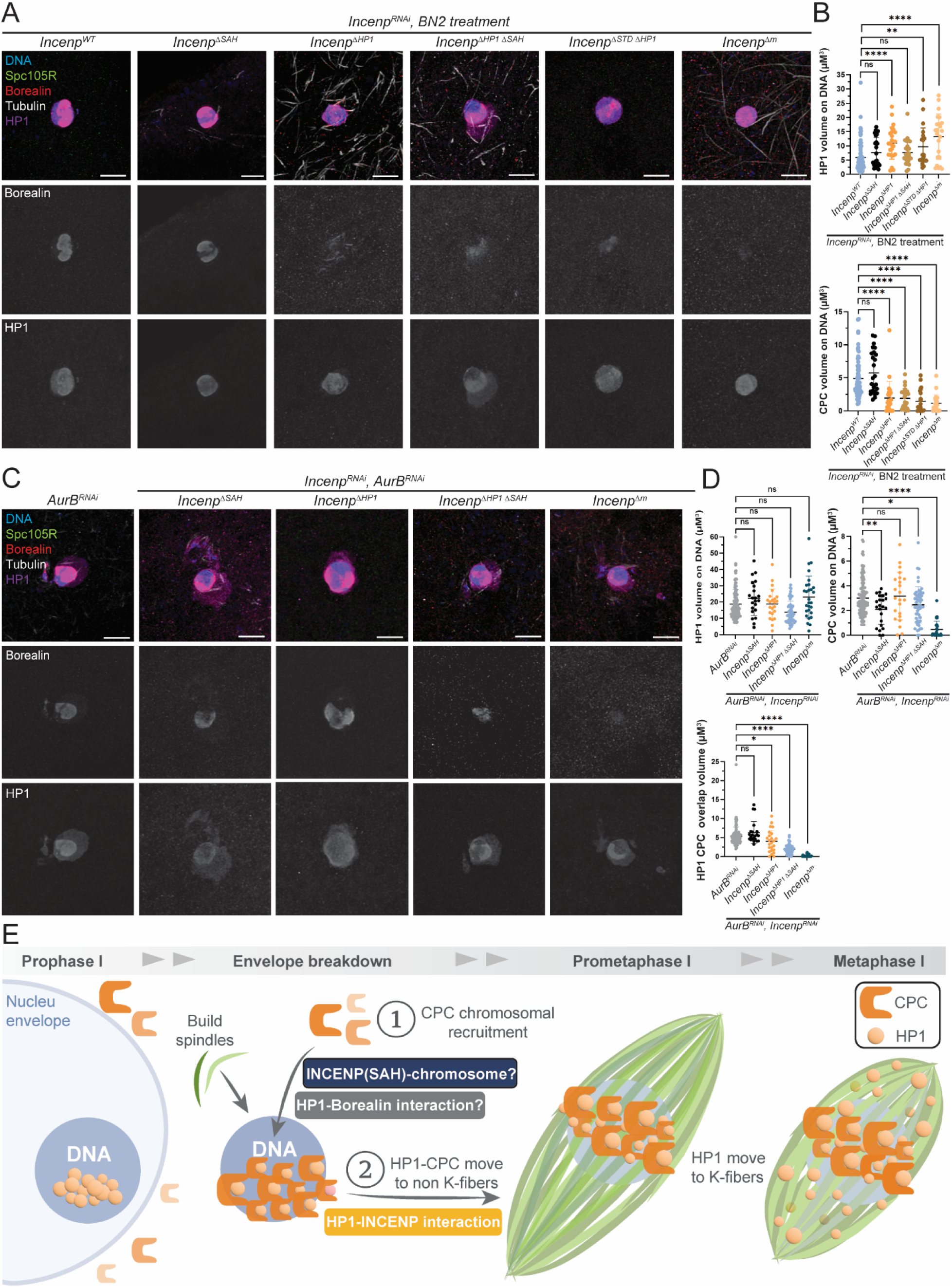
Chromosomal recruitment of the CPC does not depend on the INCENP HP1 domain. Two experiments were performed to investigate the effect of *Incenp* mutants on recruiting the CPC to the chromosomes. **(A)** BN2 treatment inhibits AurB and depolymerizes all microtubules in oocytes. In controls, this results in the return of HP1 (purple) and the CPC/Borealin (red) to the chromosomes (blue), with tubulin (white) and SPC105R (green). **(B)** Quantitative analysis of HP1 and CPC volume on chromosomes. Data was analyzed in *Incenp^WT^* (n = 78), *Incenp*^Δ*SAH*^ (n = 26), *Incenp*^Δ*HP1ex*^ (n = 25), *Incenp*^Δ*HP1*^ ^Δ*SAH*^ (n = 25), *Incenp*^Δ*STD*^ ^Δ*HP1*^ (n = 21), and *Incenp*^Δ*m*^ (n = 23) oocytes, all in a *Incenp^RNAi^* background. **(C)** Spindle assembly is prevented in *AurB^RNAi^* oocytes, and HP1 and the CPC remain on the chromosomes. **(D)** Quantitative analysis of HP1 and CPC volume on the chromosomes. HP1–CPC interaction was further assessed by measuring HP1–CPC overlap volume (lower graph). Data analysis was based on *Incenp^WT^* (n = 113), *Incenp*^Δ*SAH*^ (n = 22), *Incenp*^Δ*HP1ex*^ (n = 23), *Incenp*^Δ*HP1*^ ^Δ*SAH*^(n = 54), and *Incenp*^Δ*m*^ (n = 28) oocytes, all in a *Incenp^RNAi^* background. Error bars represent 95% confidence intervals and statistical significance was assessed by t-test: * = P < 0.05, ** = P < 0.01, **** = P <0.0001. Scale bar, 5 µm in all images. **(E)** Schematic diagram illustrating the proposed role of HP1 and INCENP in *Drosophila* female meiosis.

We analyzed *Incenp* mutants in *AurB^RNAi^* to examine CPC recruitment to the chromosomes prior to spindle assembly. Under these conditions, HP1 volume on the chromosomes remained unchanged across all *Incenp* mutants, consistent with prior evidence that INCENP is not required for HP1 chromosome localization. CPC localization was unaffected in the *Incenp*^Δ*HP1ex*^mutant. Interestingly, although *Incenp*^Δ*HP1ex*^showed no change in HP1 and CPC volume on the chromosomes, we observed a reduced overlap volume of HP1 and CPC (Figure 7D), indicating that the HP1–INCENP interaction was disrupted in this mutant. CPC localization was significantly reduced in *Incenp*^Δ*SAH*^, *Incenp*^Δ*HP1*Δ*SAH*^, and *Incenp*^Δ*m*^ oocytes (Figure 7C-D). Notably, oocytes with the *Incenp*^Δ*HP1*Δ*SAH*^ deletion showed a similar reduction of CPC volume on chromosomes as the *Incenp*^Δ*SAH*^mutant (Figure 7D). These results suggest that the SAH domain, but not the HP1-binding domain, has a role in CPC recruitment prior to spindle assembly.

In both chromosome binding assays, we observed HP1 concentrates or aggregation around the karyosome and not directly colocalizing with it (Figure 7A, C). As described above, this distribution pattern may result from HP1 phase separation occurring when INCENP interactions with the spindle are disrupted.

Differences between the two assays probably reflect whether the CPC moved to the microtubules prior to spindle disassembly. The reduction of CPC on the chromosomes in *Incenp*^Δ*HP1ex*^ mutants following BN2 but not in *AurB^RNAi^* suggests the CPC on the microtubules can return to the chromosomes in the BN2 experiment. The *AurB^RNAi^* results reveal a role for the SAH domain of INCENP in CPC recruitment to chromosomes prior to spindle assembly. This is not as severe as *Incenp*^Δ*m*^, indicating other domains contribute to initial recruitment of the CPC to the chromosomes. Because CPC chromosome recruitment was not affected in *AurB^RNAi^ Incenp*^Δ*HP1*Δ*SAH*^ oocytes, the PRD and SAH domains may be sufficient for chromosome recruitment. The failure to form spindles in *Incenp*^Δ*HP1*Δ*SAH*^ oocytes even though they showed some chromosomal CPC in the *AurB^RNAi^* assay, suggests the HP1-INCENP interaction plays a role in CPC translocation from the chromosomes to the spindle. Overall, these findings reveal distinct yet synergistic functions for the SAH and HP1 domains of INCENP in regulating CPC localization and spindle assembly.

## Discussion

### Initiation of acentrosomal spindle assembly with an HP1-CPC interaction

In this study, we conducted an analysis of the role played by chromosomal protein HP1 in oocyte meiotic spindle assembly. We found that HP1 is required to recruit the CPC to the chromosomes to initiate spindle assembly. HP1 then dissociates from the chromosomes in prometaphase I and dynamically localizes to the spindle microtubules. Our findings support a model in which HP1 plays multiple roles in *Drosophila* oocyte meiosis, starting with recruitment of the CPC, and continuing with functions in spindle assembly and function (Figure 7E). Following nuclear envelope breakdown, CPC components INCENP, Borealin, and Survivin, are recruited to the chromosomes via an interaction with HP1. Recruitment of AurB promotes the dissociation of HP1 from H3K9me3-marked chromatin by phosphorylating H3S10^52^. Dissociation of HP1 may cause movement of the CPC from the chromosomes to the microtubules. We propose that HP1 moves with the CPC to the spindle microtubules, where it contributes to spindle function and chromosome biorientation.

*Drosophila* has 5 HP1 paralogs: HP1a/Su(var)205, HP1b, HP1c, HP1d/Rhino, and HP1e. These proteins are classified based on their conserved structure, which includes an N-terminal chromodomain and a C-terminal chromoshadow domain involved in chromatin binding and protein-protein interactions, respectively^53–56^. Among these, HP1a, HP1b, and HP1c are broadly expressed in female flies, while HP1d/Rhino is predominantly expressed in the ovary, and HP1e is testis-specific^53^. These paralogs exhibit distinct subnuclear localization patterns and likely carry out specialized functions in chromatin regulation. In oocytes, HP1a is primarily associated with pericentromeric heterochromatin when cells are not undergoing cell division. HP1b localizes to both heterochromatic and euchromatic regions, whereas HP1c is mainly euchromatic^54^. HP1d/Rhino also localizes to heterochromatin but displays a pattern distinct from HP1a and the H3K9me2 chromatin mark^53^. Importantly, we have not ruled out that an HP1 paralog contributes to meiotic spindle assembly.

The HP1 interaction may create the enrichment of CPC on the chromosomes necessary for activation of AurB. We observed evidence that HP1 and the CPC can form extra-chromosomal concentrates when spindle assembly was prevented. This activity could be related to observations that HP1 and the CPC can phase separate and form condensates^26,49,57–59^. Furthermore, artificial dimerization has been shown to activate AurB kinase^60,61^. Thus, the formation of LLPS concentrates may be required for AurB to activate spindle assembly factors. Similarly, a phase separated structure may concentrate multiple microtubule-organizing factors to promote acentrosomal spindle assembly in mammalian oocytes^62^.

The CPC could promote spindle assembly by directly interacting with microtubules^46^. In support of this conclusion, the CPC localizes to all spindle microtubules in the absence of Subito, which transports it to the central spindle^39,50^. An *in vitro* study revealed that concentrated CPC purified from *Escherichia coli* can assemble parallel microtubule bundles^58^. However, there is also evidence of indirect roles. Phosphorylation by the CPC promotes kinetochore assembly, and may recruit the kinesin Subito that bundles antiparallel microtubules^63–65^. In *Drosophila* oocytes, the CPC disrupts an inhibitory interaction between 14-3-3 and spindle assembly factors, including Borealin and the kinesin-14 is NCD^66,67^. In all these examples, spindle assembly may only occur around the chromosomes because activation of AurB activity requires concentrating the CPC, which depends on interactions with chromosome-bound HP1.

### The CPC chromosomal recruitment mechanism

Our *AurB^RNAi^* data indicates that HP1 recruits the CPC for spindle assembly. In prophase, the CPC is outside and HP1 is inside the nucleus. Thus, NEB allows the CPC to interact with HP1. In the *AurB^RNAi^* chromosome binding assay, we observed that CPC localization on chromosomes was reduced to the same degree in *Incenp*^Δ*SAH*^and *Incenp*^Δ*HP1*Δ*SAH*^ oocytes (Figure 7C-D), suggesting that the SAH domain but not the HP1-binding domain, functions in the chromosomal recruitment of the CPC. This finding is consistent with previous reports in mitotic systems. For example, the SAH domain of INCENP contributes not only to microtubule binding but also supports CPC localization to centromeric chromatin in human cells and Xenopus egg extracts^51^.

*Incenp*^Δ*m*^ oocytes have a more severe phenotype than *Incenp*^Δ*SAH*^, suggesting additional domains must cooperate with SAH in recruitment to the chromosomes. Furthermore, the *Incenp*^Δ*STD*Δ*HP1*^ mutant protein is recruited to the chromosomes and forms meiotic spindles, indicating that the PRD and SAH domains are sufficient for spindle assembly. This is consistent with previous studies suggesting that the PRD domain cooperates with the SAH domain to promote CPC localization to chromatin and microtubules^51^. Whether the SAH and PRD domains interact with HP1 is not known. However, the observation that *AurB^RNAi^*, *Hp1^RNAi^* oocytes lack CPC on the chromosomes suggests the SAH interaction does depend on HP1, although it might not be direct (Figure 3C).

Co-IP assays revealed that HP1 interacts with CPC, which could include interactions INCENP and Borealin (Figure 4B). Borealin also contains an HP1-binding domain in its C-terminal domain^68^. In our previous study showed, we provided evidence that a *Borealin* - *Incenp* fusion (*borr* : *Incenp*) formed bipolar spindles that depended on the Borealin C-terminal domain^19^. This result suggests that two HP1 interactions with the CPC, HP1-Borealin and HP1-INCENP, are required for initiating spindle assembly. It is possible that a direct Borealin-HP1 interaction is required for CPC recruitment to the chromosomes, and the PRD-SAH dependence on HP1 is indirect. The importance of regulating this interaction was recently demonstrated by the analysis of *pal* mutants in *Drosophila* males^69^. The Pal protein is required for the removal of histones H3 and H4 from sperm chromatin. In *pal* mutants, the paternal chromosomes behave just like the maternal chromosomes; they recruit the CPC, have H3S10ph, and build AurB kinase-dependent spindles. It is not known if the CPC recruitment to paternal chromosomes depends on HP1.

### Does the CPC recruit HP1?

CPC levels on chromosomes following BN2 treatment and spindle depolymerization depended on the INCENP HP1 domain but not the SAH domain (Figure 7). This is different than the observations with *AurB^RNAi^* and we speculate that this difference is because returning to the chromosomes requires an HP1 – INCENP interaction first, whereas initial recruitment (where HP1 starts on the chromosomes) does not. This may be related to observations in cancer cells that HP1 binds directly to INCENP when the CPC is localized on kinetochores^70^. Similarly, we can’t rule out the possibility that during spindle assembly, the CPC localized on spindles recruits cytoplasmic HP1. Following BN2-induced spindle disassembly, this spindle-associated HP1 could return to the chromosomes.

### The mechanism of HP1-CPC movement to the spindle

Using live imaging, we directly observed the movement of HP1 off the chromosomes (Figure 1C). These observations showed that only a small amount of the chromosomal HP1 ends up on the spindle. The ejection of HP1 from the chromosomes is AurB-dependent and may result from phosphorylation of H3S10, which can inhibit the H3K9me-HP1 interaction^52,71^. Live imaging also showed there is a delay where spindle formation has begun, but HP1 remains on the chromosomes. A similar observation has been made with live imaging of the CPC ^72^. Thus, HP1 and the CPC may move together after the initiation of spindle assembly.

Four additional observations suggest that HP1 has an important function while localized to microtubules. First, the HP1 domain of INCENP is required for spindle assembly but is not required for recruiting the CPC to the chromosomes. Second, the most severe defects in biorientation were found in the *Incenp*^Δ*HP1*^ mutant. Third, the kinesin 6 Subito has an HP1 interaction site, which is required for localization of this motor to the spindle^19^. Fourth, the chromosome-independent spindle assembly phenotype observed in *subito*^Δ*N*^ mutants depends on HP1. Thus, we propose that HP1 is required for the translocation of the CPC from the chromosomes to the microtubules and is required on the spindle.

The *Incenp*^Δ*STD*Δ*HP1*^ mutant forms a bipolar spindle but has biorientation defects, indicating that the STD domain is required for spindle function. Like the chromosome binding activity, there appears to be multiple domains within the CPC that contribute to the spindle function. We propose that the HP1, SAH and STD domains within INCENP and a domain within Borealin are required for spindle-based interactions and functions^19^. An association with the spindle has been suggested to be required for the CPC to destabilize incorrectly attached kinetochores^62^. A similar activity has been described in mitotic anaphase of *Drosophila* and other organisms, where CPC in the midzone can detect lagging chromosomes^73–75^ and an AURB phospho-gradient generated from the midzone MTs can destabilize incorrect attachments^76,77^. We suggest that CPC localized to the central spindle of *Drosophila* metaphase I oocytes destabilizes KT-MT attachments. Similarly during prometaphase in mouse oocytes, the chromosomes congress to the central spindle^78^ and KT-MT attachments, regardless of biorientation, are destabilized because of high AURB/AURC activity^79^. These results suggest that spindle-localized CPC that regulates kinetochore – microtubule attachments and functions in error correction may be a conserved in mammalian and *Drosophila* oocytes.

## Materials and Methods

### Drosophila genetics, generation of RNAi-resistant transgenes and shRNAs

All *Drosophila melanogaster* strains were crossed and maintained on BDSC Standard Cornmeal Medium (https://bdsc.indiana.edu/information/recipes/bloomfood.html) at 25°C. Fly stocks were obtained from the Bloomington Drosophila Stock Center including *aurB* (*GL00202*) and *Incenp* (*GL00279*). Males carrying an shRNA regulated by the UAS promoter were crossed to *mata4-GAL-VP16* to achieve post-pachytene expression in the female germline ^80^. Two shRNAs targeting *Su(var)205 (HP1)* were designed and cloned into pVALIUM22 according to the protocols at the Transgenic RNAi Project (TRiP) at Harvard Medical School and injected into *Drosophila* embryos by Model System Injections or BestGene. For live imaging, a GFP fusion at the endogenous locus (*TI{TI}Su(var)205^EGF^*^P^) ^81^ and *ubiq:: Jupiter*.*mCherry-* ^82^ were used. For co-IP experiments, we used HA fusions to *HP1* ^83^ and *aurB* under the control of UASp. The *aurB* transgene was constructed fusing the coding region in cDNA LD39409 to the 3XHA tag in pPHW.

The *Incenp* mutants used in this work, the deleted regions, and their fertility are summarized in Table 1. *Incenp* transgenes were generated using the fully sequenced *Incenp* cDNA RE52507 obtained from the Drosophila Genomic Resource Center. The cDNA was subcloned into the pENTR4 vector (Invitrogen), and eight silent mutations were introduced at the site targeted by the *Incenp* shRNA (*GL00279*), which corresponds to the region encoding amino acids 437–441 ^19^. This wild-type *Incenp* cDNA was cloned into a UASP vector. All mutant transgenes were constructed using this *wild-type Incenp* as the backbone (Genscript). We previously generated an *Incenp* mutant with a deletion of the HP1-binding region spanning amino acids 121–232 in *Drosophila*^19^. However, the HP1 interaction region of INCENP may be larger than expected ^29^. To ensure complete removal of all potential HP1-binding sites, we generated a new *Incenp*^Δ*HP1ex*^mutant by deleting 121–397 amino acids.

Each *Incenp* deletion mutant was injected into embryos by Model System Injections (Durham, NC) or BestGene. For each experiment, males carrying a *Incenp* RNAi resistant transgene and *GL00279* were crossed to the *mata4-GAL-VP16* (*matα-Gal4*), inducing both shRNA and RNAi-resistant transgene expression after pachytene and throughout all stages of prophase in oocyte development^80,84^. At least two independent transgenic lines of each construct were tested for their ability to rescue *Incenp^RNAi^* using shRNA GL00279. The generation and genetic analysis of *GFP-sub*^Δ*N*^ construct were previously described^50^.

### Nondisjunction and female fertility assays

To assess chromosome nondisjunction and female fertility, virgin females of each genotype were crossed to *y w/B^S^Y* males. If nondisjunction occurred, females would be observed with a bar-eyed phenotype (XXY), and males with a normal non-bar-eyed phenotype. The XXX and OY genotypes are lethal, which is accounted for in the following equation used to calculate the rate of nondisjunction:# NDJ flies/ [(#nondisjunction fliesX2)+(normal flies)]. Progeny were counted 18 days after each cross. Female fertility was quantified by dividing the total number of adult progenies per vial by the number of parental females.

### Prophase oocyte preparation for immunofluorescence

Whole ovaries were prepared for immunofluorescence as previously described^85,86^. Briefly, ovaries were dissected from approximately 20 mated females aged 1–2 days in 1× Modified Robb’s Buffer. Within 20 minutes, the dissected ovaries were fixed in a buffer containing 4% paraformaldehyde for 9 minutes at room temperature, followed by washes in BAT buffer (1.5 mM PIPES, 80 mM KCl, 20 mM NaCl, 2mM EDTA, 0.5mM EGTA, 0.1% TritonX, 1mM DTT, 0.5 mM Spermidine, 0.15 mM Spermine). BAT buffer supplemented with normal goat serum (BAT-NGS, 6 mg/ml) was used to block nonspecific antibody binding. Samples were then incubated with primary antibodies overnight at 4°C (Supplementary Table 1). The following day, tissues were washed and incubated with secondary antibodies that had been preabsorbed with fixed *Drosophila* embryos. DNA was stained using Hoechst 33342 (10 μg/ml). Final washes were performed in BAT buffer followed by 1× Buffer A, and samples were mounted in SlowFade Gold Antifade (Invitrogen) for imaging.

### Stage 14 (Prometaphase I - Metaphase I) oocyte preparation for immunofluorescence

The collection, preparation, and imaging protocol of mature *Drosophila* oocytes were performed as previously described^87^. Briefly, stage 13–14 oocytes were collected in 1× Modified Robb’s Buffer (55mM sodium acetate (NaOAc), 40mM potassium acetate (KOAc), 100mM sucrose, 10mM dextrose, 1.2mM MgCl_2_, 1mM CaCl_2_, 100mM HEPES, PH 7.4) from approximately 200 two-day-old, yeast-fed, non-virgin females raised at 25 °C. Oocytes were fixed in 4% formaldehyde buffer (1 x PBS, 105 mM sucrose) for 2.5 minutes, then in heptane for 1 minute. Following fixation, oocytes were manually rolled to remove the surrounding membranes, incubated in 1 x PBS with 1% Triton X-100 to enhance permeabilization, and blocked in PTB (1 x PBS + 10% Tween-20 + 0.5% BSA). Oocytes were then incubated with primary antibodies (Supplementary Table 1), followed by fluorophore-conjugated secondary antibodies preabsorbed against embryos to reduce non-specific binding. DNA was counterstained with Hoechst 33342 (10 μg/ml). After staining, oocytes were washed in PTB and mounted in SlowFade Gold Antifade Mountant (Invitrogen) for imaging.

To inhibit Aurora B kinase activity, oocytes were incubated in 50 µM BN2 in 0.1% DMSO prepared in 1× modified Robb’s buffer for 60 minutes prior to fixation. This treatment results in spindle disassembly due to microtubule depolymerization. Each experiment was tested at least twice in independent biological replicates.

### Fluorescence in situ hybridization (FISH) of oocytes

Stage 13–14 oocytes were collected as described for immunofluorescence but fixed using a formaldehyde/cacodylate-based fixation solution^88^. The fixation buffer consisted of 600 mM sucrose, 800 mM potassium acetate (KOAc), 800 mM sodium acetate (NaOAc), and 200 mM EGTA, with the addition of 8% formaldehyde and 100 mM cacodylate acid. Oocyte Membranes were removed by rolling in 2× SSCT media, followed by graded washes in 20%, 40%, and 50% formamide. Samples were nutated in 50% formamide for 5 hours at 37 °C to enhance probe penetration. FISH probes targeting the X-chromosome 359 (Alexa Fluor 594), second chromosome AACAC (Cy3), and third chromosome dodeca (Cy5) repeats were synthesized by Integrated DNA Technologies (IDT). The probes were added to oocytes in hybridization buffer containing dextran sulfate, formamide and SSC, and incubated in a thermocycler for 20 min at 80 °C for denaturation, followed by overnight incubation at 37 °C. After hybridization, samples were transferred into 50% and then 20% formamide to gradually decrease the formamide concentration, and then blocked in PTB. Microtubules and DNA were labeled with anti-tubulin–FITC (Sigma-Aldrich; 1:30) and Hoechst 33342 (10 µg/ml), respectively. Oocytes were washed in PTB and mounted in SlowFade Gold Antifade (Invitrogen) for imaging. At least three independent biological replicates per genotype were performed to test the consistency of chromosome biorientation rates.

### Live Imaging of *Drosophila* Oocytes

Females expressing fluorescently tagged proteins of interest were maintained on yeast for 2–3 days at 25 °C prior to dissection. Ovaries were dissected in Halocarbon 700 oil (Sigma) at room temperature. Individual oocytes were gently separated using forceps and spread onto a 42 × 50 mm coverslip. For imaging meiotic spindle assembly, late stage 12 oocytes lacking dorsal appendages were selected. Chromatin and nuclear envelope dynamics were visualized using HP1-GFP, while microtubules were labeled with Jupiter-mCherry. All dissections and mounting steps were completed within 20 minutes to preserve physiological conditions. Live imaging was performed on an Andor Dragonfly Spinning Disk Confocal Microscope (Dragonfly system) under the 63×/1.4 oil-immersion objective. Prior to imaging, system alignment and connectivity were verified, and immersion oil was applied to the objective. Z-stacks were acquired with a step size of 0.8 µm. Time-lapse imaging was initiated at nuclear envelope breakdown (NEBD), where the nuclear envelope deforms and tubulin accumulates around chromatin. Acquisition frequency was set to every 3 minutes to capture spindle assembly dynamics.

### Embryo cytology

Embryos were collected from mated females of the desired genotype maintained on grape juice agar plates supplemented with yeast paste to stimulate oviposition. Collections began on day 2, and embryos were harvested in 1-hour laying intervals followed by 1 hour of aging to enrich for syncytial mitosis division stages. Embryos were dechorionated using 50% bleach for 2 minutes, rinsed thoroughly with distilled water, and transferred to a 1:1 mixture of heptane and methanol for 30 seconds with vigorous shaking to remove the vitelline membrane. Following membrane removal, embryos were fixed overnight at 4°C in methanol. Fixed embryos were gradually rehydrated through a methanol/PBS series (80:20, 60:40, and 20:80), washed in PBS, and blocked in PTB for 1 hour at room temperature. Primary antibodies were applied in PTB and incubated overnight at 4°C. Following washes in PTB, embryos were incubated with fluorophore-conjugated secondary antibodies and Hoechst 33342 (10 μg/ml) for nuclear staining. After final washes, embryos were mounted on glass slides for imaging.

### RNA Extraction and RT-PCR

To quantify the efficiency of HP1 knockdown by RNAi, total RNA was isolated from oocytes using TRIzol Reagent (Thermo Scientific) according to the manufacturer’s protocol. cDNA was synthesized from 2 µg total RNA from each sample using the High-Capacity cDNA Reverse Transcription Kit (Applied Biosystems). Quantitative RT-PCR was performed using TaqMan Gene Expression Assays (Thermo Scientific) on a QuantStudio Real-Time PCR System. Relative gene expression levels were calculated using the ΔΔCt method and normalized to internal controls.

### Western blotting and co-immunoprecipitation

Western blotting was performed as previously described^37^. Oocytes were isolated from yeasted females in phosphate-buffered saline (PBS) and then ground and boiled in 2× Laemmli Sample Buffer supplemented with protease inhibitor cocktail (Roche). 1 mg of total protein was loaded per lane on SDS-PAGE gels and transferred to a nitrocellulose membrane (Bio-Rad) using a Trans-Blot Turbo machine (Bio-Rad). Membranes were blocked with Intercept Blocking Buffer (LI-COR) for 1h at room temperature, then sequentially incubated with primary and secondary antibodies from LI-COR (Supplementary Table 1). Blots were imaged using the Odyssey CLX Imaging System (LI-COR Biosciences).

Co-immunoprecipitation was performed as previously described^89^. The *aurB* and *Hp1* transgenes were constructed to include three tandem HA epitope tags, and because they were under the control of the UAS promoter, were expressed using *mata4-GAL-VP16*. Protein extracts were prepared from embryos lysed in IP buffer containing 50 mM Tris-HCl (pH 8.0), 400 mM NaCl, 0.5% (v/v) NP-40, 0.1% (w/v) sodium deoxycholate, and a protease inhibitor cocktail. A 25 μL aliquot of the lysate was saved as input. The remaining lysate was then incubated overnight at 4°C with 20 μL of anti-HA agarose beads (HA-Tag IP/Co-IP Application Set, Thermo Scientific, Cat. #26180Y). Following incubation, beads were washed with Tris-buffered saline containing 0.05% Tween-20 (TBST; 25 mM Tris-HCl, 150 mM NaCl, 0.05% Tween-20; pH 7.2). Bound proteins were eluted by adding 2× Non-Reducing Sample Buffer directly to the beads. Elution and input samples were analyzed by SDS-PAGE. At least three independent biological replicates were performed to exclude sample variation.

### Cytology and fluorescent image analysis

Imaging was performed on a Leica TCS Stellaris 8 confocal microscope equipped with a 63X, N.A. 1.4 oil immersion objective. Images stacks of germariums, oocytes, and embryos were acquired in XYZ dimensions and shown as maximum intensity projections. Cropping was performed in Adobe Photoshop CS6 (Adobe, San Jose, CA). Stage 14 oocyte images were analyzed using Imaris image analysis software (Bitplane), following the parameters described in Wang et al^19^. Colocalization and volume measurements were quantified based on distinct fluorescence channels in individual oocytes. Images were used to quantify the volumes and fluorescence intensities of CPC, HP1, and microtubules. To assess colocalization of CPC and HP1 with microtubules and DNA, surface objects corresponding to the spindle, DNA, HP1, and CPC were generated. Colocalization was quantified by measuring the overlap volume between the corresponding surface objects. Quantifications were based on at least 18 oocytes from three independent biological replicates.

Chromosome biorientation was measured in FISH assays. In FISH images, each data point represents one homologous centromere pair. Homologs were scored as bi-oriented if the two FISH signals were located on opposite sides of the chromatin mass. Homolog pairs with both signals on the same side of the spindle were considered defective. At least four independent biological replicates per genotype were performed to determine the consistent phenotype.

### Statistical Analysis

Statistical analyses were performed using Prism software (GraphPad), with tests specified in individual figure legends. Tests included unpaired two-tailed Student’s *t*-test for comparisons between two groups, one-way ANOVA for comparisons among multiple groups, and Fisher’s exact test for categorical data. Data are presented as mean ± standard deviation (SD) from *n* = 2–5 independent experiments. A *p*-value < 0.05 was considered statistically significant.

## Acknowledgements

Some stocks used in this study were obtained from the Bloomington Drosophila Stock Center (Bloomington, IN). We thank Christian Lehner (University of Zurich, Zurich, Switzerland) for generously providing antibodies. We also acknowledge the Waksman Institute Shared Imaging Facility for access to confocal microscopy equipment and technical support. Siwen Wu was supported by a Busch Predoctoral Fellowship. This work was funded by the National Institutes of Health (grant GM101955 to K.S.M.). The authors declare no competing financial interests.

