## Supplemental Material for "An interaction between HP1 and the Chromosomal Passenger Complex Initiates Acentrosomal Spindle Assembly in *Drosophila* oocytes"

### Supplemental Tables and Figures

#### Supplementary Table 1: Antibodies used in this work

| Antibody (RRID) | Host species | Vendor or reference | Catalog No. | Application (working dilution) |
|---|---|---|---|---|
| Borealin | Guinea pig | [90] | N/A | IF (1:2,000)<br>WB (1:5,000) |
| C(2)M | Rabbit | [91] | N/A | IF (1:400) |
| CENP-C | Guinea pig | [92] | N/A | IF (1:1,000) |
| GFP (AB_2536526) | Rabbit | Thermo Scientific | G10362 | IF (1:200) |
| HA (AB_390919) | Rat | Roche | 11867431001 | IF (1:50)<br>WB (1:2,500) |
| HP1 (AB_528276) | Mouse | DSHB | c1a9 | IF (1:50)<br>WB (1:2,500) |
| INCENP | Rat | [93] | N/A | IF (1:400)<br>WB (1:2,500) |
| SPC105R | Rabbit | [94] | N/A | IF (1:4,000) |
| Subito | Rat | [37] | N/A | IF (1:150) |
| Tubulin (AB_528499) | Mouse | DSHB | E7 | IF (1:250) |
| Tubulin (AB_93477) | Rat | Millipore | CBL270-I | IF (1:300) |
| Tubulin (AB_477583) | Mouse | Sigma-Aldrich | T6199 | WB (1:4,000) |
| Tubulin-FITC (AB_476967) | Mouse | Sigma-Aldrich | F2168 | FISH (1:30) |
| Alexa Fluor 488 anti-Mouse IgG (AB_2534088) | Goat | Thermo Scientific | A-11029 | IF (1:75) |
| Alexa Fluor 488 anti-Rabbit IgG (AB_2313584) | Donkey | Jackson ImmunoResearch | 711-545-152 | IF (1:200) |
| Alexa Fluor 594 anti-Mouse IgG (AB_2340854) | Donkey | Jackson ImmunoResearch | 715-585-150 | IF (1:200) |
| Alexa Fluor 594 anti-Rabbit IgG (AB_2534095) | Goat | Thermo Scientific | A-11037 | IF (1:100) |
| Alexa Fluor 647 anti-Guinea Pig IgG (AB_2340476) | Donkey | Jackson ImmunoResearch | 706-605-148 | IF (1:100) |
| Alexa Fluor 647 anti-Rat IgG (AB_2340694) | Donkey | Jackson ImmunoResearch | 712-605-153 | IF (1:300) |
| Cy3 anti-Guinea Pig IgG (AB_2340460) | Donkey | Jackson ImmunoResearch | 706-165-148 | IF (1:200) |
| Cy3 anti-Rabbit IgG (AB_2338006) | Goat | Jackson ImmunoResearch | 111-165-144 | IF (1:250) |

| | | | | |
|---|---|---|---|---|
| Cy3 anti-Rat IgG (AB_2338251) | Goat | Jackson ImmunoResearch | 112-165-167 | IF (1:200) |
| IRDye 680RD anti-Mouse IgG (AB_10956588) | Goat | LICORbio | 926-68070 | WB (1:20,000) |
| IRDye 680LT anti-Rat IgG (AB_10715073) | Goat | LICORbio | 926-68029 | WB (1:20,000) |
| IRDye 680RD anti-Guinea Pig IgG (AB_10956079) | Donkey | LICORbio | 926-68077 | WB (1:20,000) |
| Alexa Fluor 680 anti-Guinea Pig IgG (AB_2340478) | Donkey | Jackson ImmunoResearch | 706-625-148 | WB (1:20,000) |
| IRDye 800CW anti-Mouse IgG (AB_2687825) | Goat | LICORbio | 925-32210 | WB (1:20,000) |
| IRDye 800CW anti-Guinea Pig IgG (AB_1850024) | Donkey | LICORbio | 926-32411 | WB (1:20,000) |

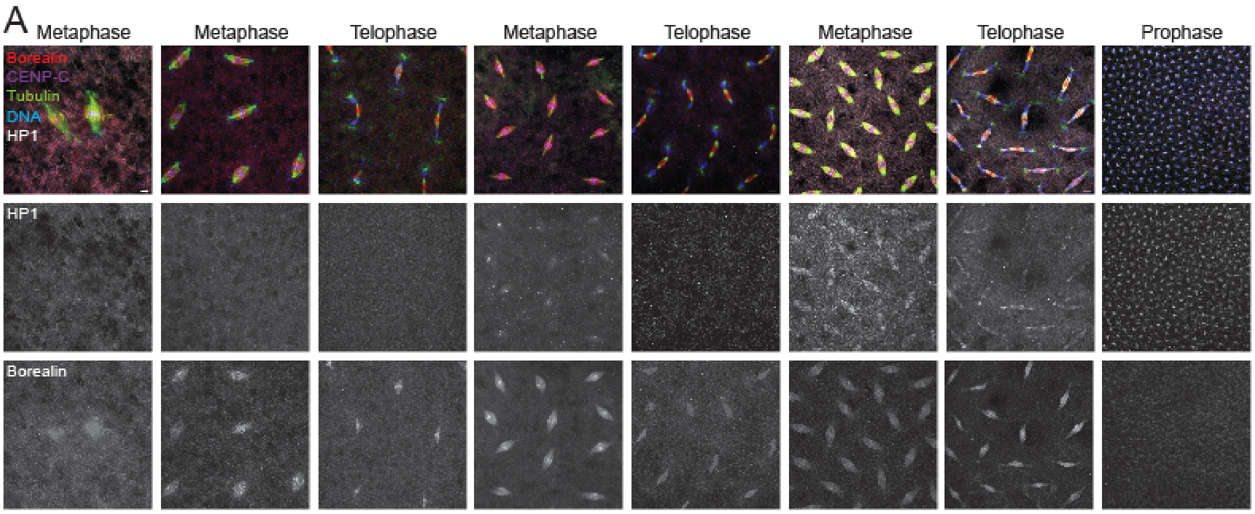


Supplemental Figure 1. **HP1 localization pattern during mitosis in embryos and in the ovary after knockdown.**

**(A)** *Drosophila* embryos from wild-type females were collected at 2-hour intervals. Borealin is shown in red, DNA in blue, HP1 in white, CENP-C in purple, and microtubules in green. The HP1 and Borealin channels are also shown separately. Scale bar is 5 μm. HP1 is absent during early stages of embryonic development but appears at later mitotic stages. HP1 is found on the chromosomes in prophase and relocates to the spindle microtubules at metaphase. Scale bar is 5 μm.

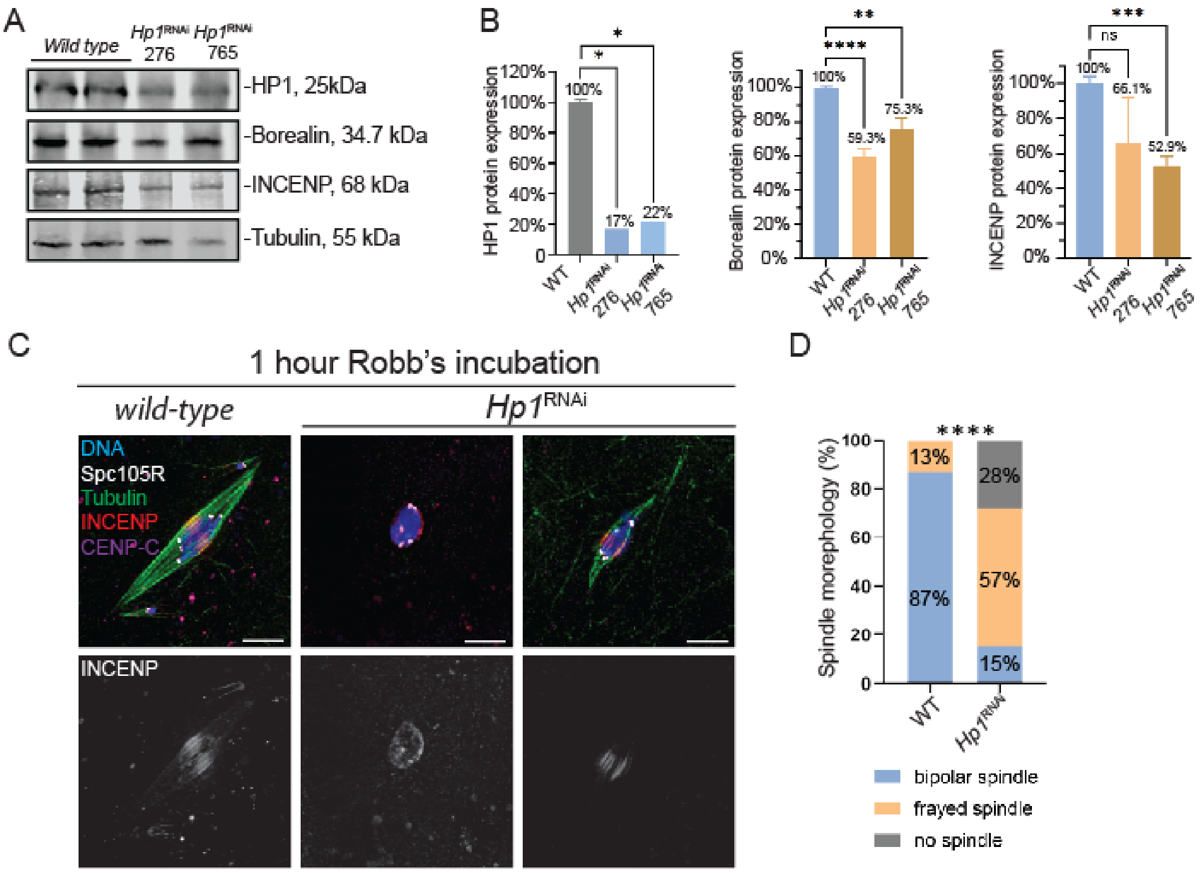


Supplemental Figure 2. **HP1 knockdown and effect on spindle stability.**

**(A)** Immunoblot of HP1 and CPC components INCENP and Borealin expression level following HP1 knockdown (KD) by two RNAi lines, with Tubulin serving as a loading control. Each lane contained 1 mg of total protein. **(B)** Bar graphs showing knockdown of HP1 protein (~80% reduction) and Borealin and INCENP proteins by expression of HP1 shRNAs. (* $P < 0.05$, ** $P < 0.01$, *** = p-values < 0.001, **** = p-values < 0.0001, and ns = no significance with unpaired two-tailed t-test). **(C)** To test spindle stability, both *wild-type* and *Hp1*$^{RNAi}$ oocytes were incubated in modified Robb's buffer for an hour and analyzed for changes in spindle morphology. Representative images are shown. DNA is shown in blue, SPC105R in white, Tubulin in green, INCENP in red, and CENP-C in purple. Scale bar is 5 µm. **(D)** Quantification of spindle defects from 35 *wild-type* and 73 *Hp1*$^{RNAi}$ oocytes, collected across four independent biological replicates. Quantification of spindle defects was analyzed using Fisher's exact test. Statistical significance: **** P < 0.0001.

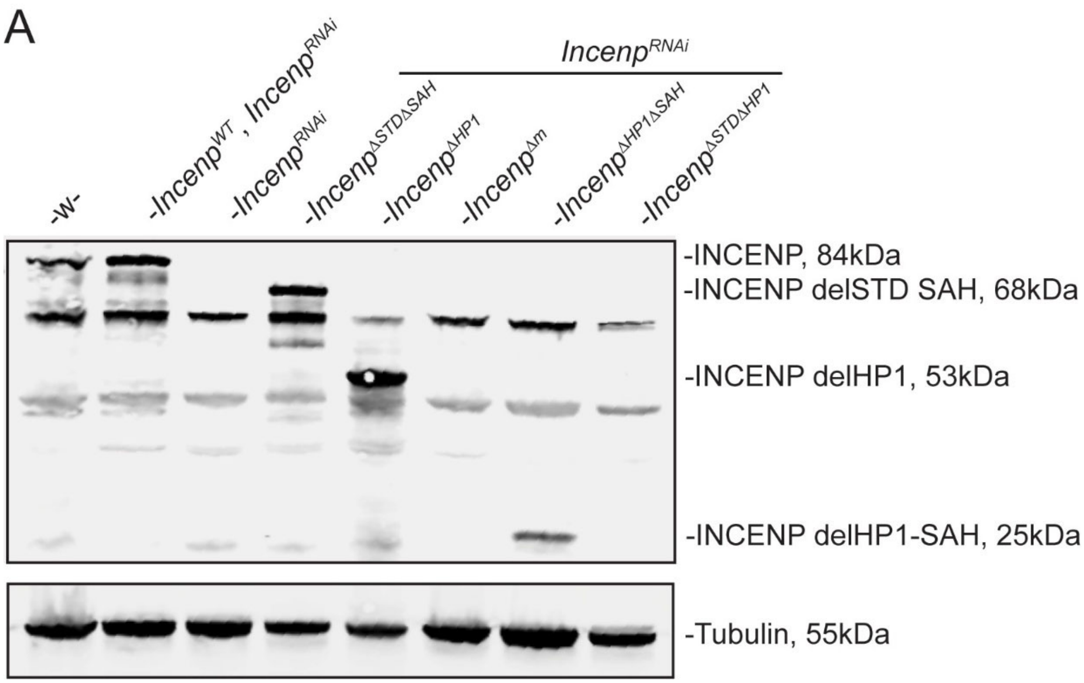


Supplemental Figure 3. **Expression of *Incenp* mutant transgenes in oocytes.**

**(A)** Immunoblot showing the expression of *Incenp* mutant proteins in oocytes in a background of *Incenp* RNAi. Tubulin was used as a loading control. Each lane was loaded with 1 mg of total protein.

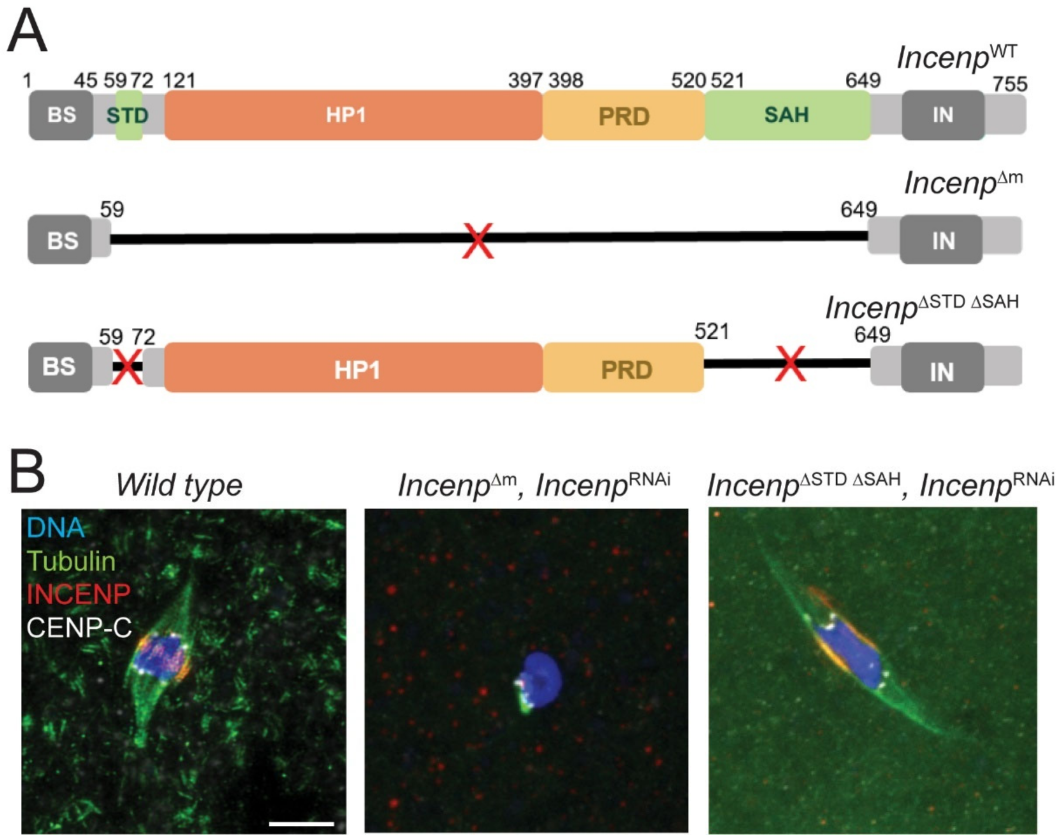


Supplemental Figure 4. **MT-binding domains of INCENP are not essential for spindle assembly or CPC localization in *Drosophila* oocytes.**

**(A)** Schematic of *Drosophila* wild-type and two mutant INCENP transgenes, $Incenp^{\Delta m}$ , which deletes all 4 domains, and $Incenp^{\Delta STD \Delta SAH}$, which deletes the proposed microtubule binding domains. **(B)** Immunofluorescence of $Incenp^{\Delta m}$ and $Incenp^{\Delta STD \Delta SAH}$ stage 14 oocytes, with tubulin (green) and CPC/INCENP (red), and CENP-C (white). Scale bar is 5 µm.

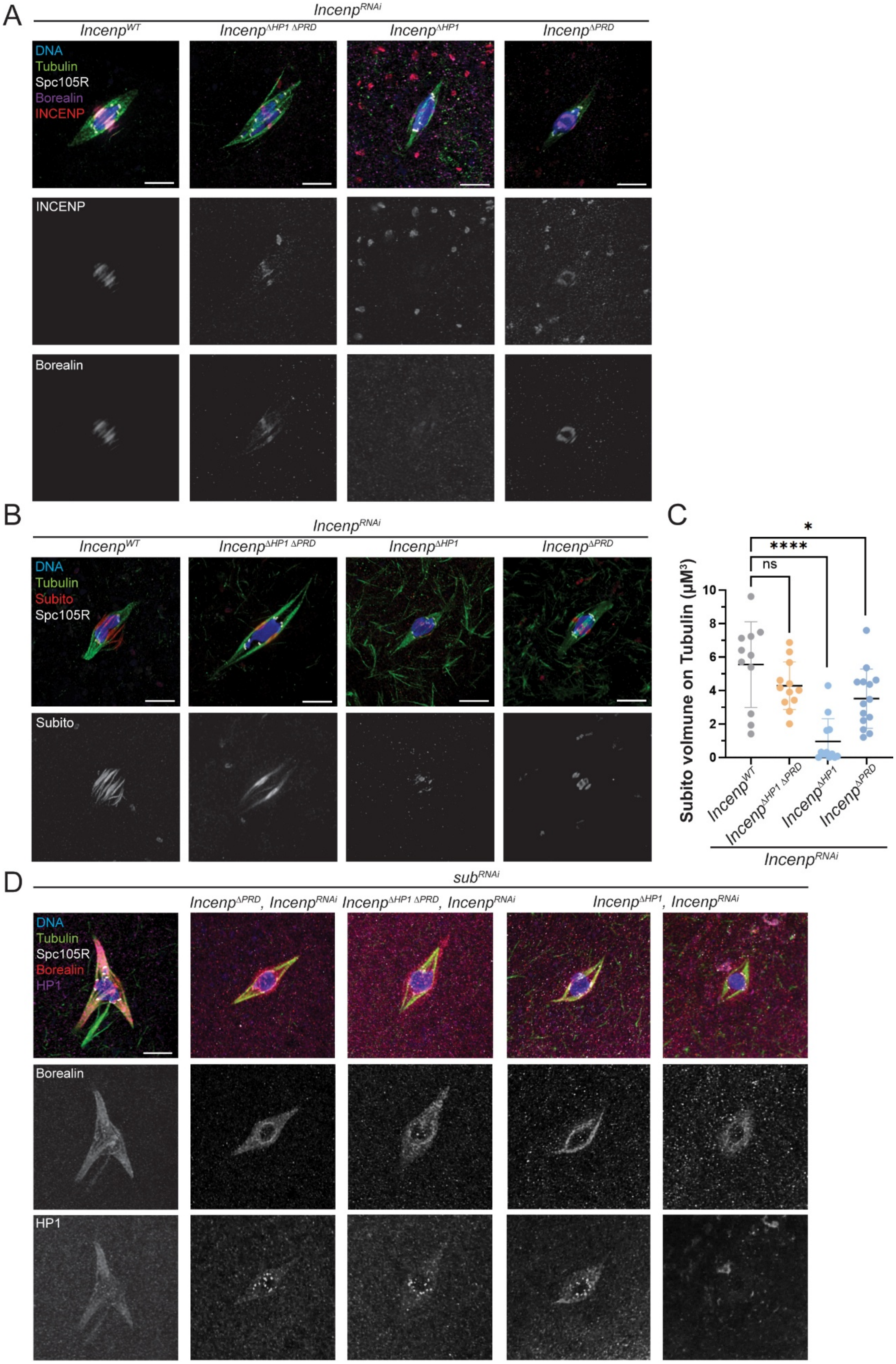
A
$Incenp^{RNAi}$
$Incenp^{WT}$
$Incenp^{\Delta HP1\ \Delta PRD}$
$Incenp^{\Delta HP1}$
$Incenp^{\Delta PRD}$
DNA
Tubulin
Spc105R
Borealin
INCENP
INCENP
Borealin
B
$Incenp^{RNAi}$
$Incenp^{WT}$
$Incenp^{\Delta HP1\ \Delta PRD}$
$Incenp^{\Delta HP1}$
$Incenp^{\Delta PRD}$
DNA
Tubulin
Subito
Spc105R
Subito
C
*
****
ns
Subito volmune on Tubulin (μM$^3$)
10
8
6
4
2
0
$Incenp^{WT}$
$Incenp^{\Delta HP1\ \Delta PRD}$
$Incenp^{\Delta HP1}$
$Incenp^{\Delta PRD}$
$Incenp^{RNAi}$
D
$sub^{RNAi}$
$Incenp^{\Delta PRD}$, $Incenp^{RNAi}$
$Incenp^{\Delta HP1\ \Delta PRD}$, $Incenp^{RNAi}$
$Incenp^{\Delta HP1}$, $Incenp^{RNAi}$
DNA
Tubulin
Spc105R
Borealin
HP1
Borealin
HP1

Supplemental Figure 5. **Disruption of the HP1–INCENP interaction leads to CPC mislocalization and central spindle defects.**

**(A)** *Incenp* mutants deleting the HP1-binding and PRD domains were stained for INCENP (red), Borealin (purple), tubulin (green) and SPC105R (white) to assess the localization of CPC. Representative images are shown. Subito knockdown causes CPC localization to all spindle microtubules. **(B)** The localization of the kinesin Subito (red) in *Incenp* mutants. **(C)** Quantification of Subito localization in *Incenp* mutant oocytes (n= 11, 12, 12 and 14). Error bars represent 95% confidence intervals. Statistical significance was assessed using Student's t-test: * P < 0.05, **** P <0.0001, ns = no significance. Scale bar is 5 μm. **(D)** *Incenp* mutant transgenes with $\textit{Subito}^{RNAi}$ in oocytes, with Borealin (red), HP1 (purple), tubulin (green), and SPC105R (white). There were two categories of $\textit{Incenp}^{\Delta HP1ex}$ oocytes. Tubulin was reduced and disorganized and HP1 was delocalized in 17 out of 30 $\textit{Incenp}^{\Delta HP1ex}$ oocytes.
